# *In vivo* crosslinking and cryo-EM analyses suggest a revised model of conjugative pilus biogenesis

**DOI:** 10.64898/2026.11.16.767151

**Authors:** Magnus Bloch, Abhinav Vadakkepat, Riccardo Zenezini, Konstantinos Thalassinos, Gabriel Waksman

## Abstract

Bacterial conjugation requires elaboration of a conjugative pilus by membrane-embedded transport machineries known as Type IV Secretion Systems (T4SSs). Recent structural findings suggested a model of conjugative pilus biogenesis in which the pilus is assembled from the VirB6-base of the T4SS Stalk, while the VirB5-apex is projected at the pilus tip. In this study, we show that *in vivo* crosslinking of these two Stalk segments within the R388 T4SS has only a modest effect on conjugative transfer efficiency, inconsistent with them separating during pilus elaboration. Combined with cryo-electron microscopy analyses of the homologous pKM101 T4SS, this leads us to propose a revised model, in which the sixfold-symmetrical inner membrane complex of the T4SS serves directly as a base for assembly of the fivefold-symmetrical pilus. The resulting symmetry mismatch is resolved by the sequential incorporation of pilus subunits at a continuously repositioning gap at the base of the nascent pilus. This revised model answers the long-standing question of how T4SSs accommodate a central symmetry mismatch during a crucial process of bacterial conjugation underpinning horizontal gene transfer.

## INTRODUCTION

Conjugation is the main pathway by which bacteria achieve horizontal gene transfer and thereby dissemination of antibiotic resistance. Conjugative plasmids encode multiprotein assemblies known as Type IV Secretion Systems (T4SSs), which function as complex nanomachines that carry out conjugative plasmid transfer. T4SSs generally consist of one VirD (VirD4) and 11 VirB (VirB1-11) proteins assembling in various stoichiometries. In this study, we refer to each VirB/D protein using the recently suggested unified T4SS nomenclature (Christie *et al*, 2026) in which the original subunit names (Trw for R388; Tra for pKM101) are followed by a subscript denoting the VirB/D subunit identity.

The T4SS encoded by the broad host range R388 plasmid consists, in its resting, pre-activation state (Macé *et al*, 2022; Macé C Waksman, 2024), of i) an outer membrane core complex (OMCC) made of 14 to 16 copies of TrwH_B7_, TrwF_B9_, and the C-terminal domain of TrwE_B10_, ii) an inner membrane complex (IMC) made of 6 copies of a protomer (hereafter referred to as the ’IMC protomer’), each containing 1 molecule of TrwM_B3_, 2 molecules of the TrwK_B4_ ATPase, an unknown number of copies of the N-terminal extension of TrwE_B10_, and 4 copies of the TrwG_B8_ C-terminal tail, iii) a periplasmic complex, the Stalk, made of a pentamer of TrwI_B6_ anchored in the inner membrane and serving as a base for a pentamer of TrwJ_B5_, and, finally, iv) another periplasmic complex, the Arches, made of 24 copies of the N-terminal domain of TrwG_B8_ and surrounding the Stalk. The TrwD_B11_ and TrwB_D4_ ATPases, along with TrwK_B4_, coordinate and power conjugation by associating with the IMC. The conjugative pilus, which is essential for conjugative substrate transfer, is assembled by the T4SS of a donor cell and extends into the extracellular environment to make direct contact with recipient cells.

Recently, we determined the high-resolution cryo-EM structure of the R388 conjugative pilus (Vadakkepat *et al*, 2024). The conjugative pilus is a 5-start helical polymer of binary complexes of the pilin protein, TrwL_B2_, and a tightly bound phospholipid (Costa *et al*, 2016). These pilus subunits form a tube, which is hypothesised to serve as a conduit for substrates during conjugative transfer (Babić *et al*, 2008; Goldlust *et al*, 2023; Beltrán *et al*, 2024). A feature of the R388 pilus structure, shared by that of the pKpQIL pilus (Zheng *et al*, 2020), is the presence of not just one but *two* helical arrangements of the pilus subunits: One apparent from the architecture of the five longitudinal strands spanning the length of the pilus, and another emerging within the resulting, transverse, pentameric rings. In the R388 pilus structure, the five subunits of each ring are related to each other by a helical twist of 66° and a slight rise of 2.6 Å, which constitutes a 1-start helical symmetry operation applied along the entire pilus. This geometric feature effectively introduces a distinct gap in each of the transverse pentamers, including the ones found at either end of the pilus (Vadakkepat *et al*, 2024). All other conjugative pilus structures reported in the literature display only 5-start helical symmetry (Costa *et al*, 2016; Amro *et al*, 2023; Kreida *et al*, 2023; Beltran *et al*, 2023; Ishimoto *et al*, 2025).

The similarities in symmetry between the R388 T4SS Stalk and the pilus led to the hypothesis that the bottom part of the Stalk, a pentamer of TrwI_B6_, serves as a base for assembly of pentameric layers of pilus subunits (Macé *et al*, 2022). Over many cycles of recruitment and translocation, a pilus growing from its TrwI_B6_ base would push the top part of the Stalk, a pentamer of TrwJ_B5_, through the OMCC, and out of the cell envelope, consistent with the known location of VirB5 at the distal end of the pilus (Aly C Baron, 2007). In this model, the two pentameric parts of the Stalk are thus completely separated, as TrwI_B6_ remains anchored to the inner membrane, while TrwJ_B5_ is pushed out. Experimental evidence indicates direct involvement of IMC ATPases in pilus subunit extraction and translocation (Kerr C Christie, 2010), while the exact mechanistic roles of the remaining subunits remain unclear. A particularly important outstanding question is how the symmetry mismatch between the pentameric Stalk/conjugative pilus and the hexameric IMC is accommodated and what role it plays during pilus biogenesis.

In this study, we present experimental evidence that necessitates revision of current models of pilus biogenesis. We use site-specific, UV-induced, *in vivo* crosslinking to demonstrate that, contrary to expectations, covalent linkage of TrwI_B6_ with TrwJ_B5_ within the R388 T4SS Stalk has only modest effects on conjugative transfer efficiency. Conflicting with prevailing models of pilus biogenesis, it thus appears that separation of the TrwJ_B5_ and TrwI_B6_ pentamers is *not* required for conjugation and, therefore, pilus elaboration. In addition, we show that inter-subunit crosslinking of the TrwI_B6_ pentamer has a strong, negative effect on R388 conjugation efficiency, indicating that this Stalk component is dynamically rearranged during the conjugation process. Finally, by solving the cryo-EM structure of the IMC protomer of the pKM101 T4SS, a close homologue of the R388 T4SS, we provide evidence that the major pilus protein, VirB2, interacts directly with the IMC component VirB3 in a manner similar to, yet different from, what is predicted by co-evolution and AlphaFold analyses.

Our experimental results lead us to propose a revised model of conjugative pilus biogenesis: VirB2 pilus subunits are recruited directly to the IMC via six symmetrically arranged copies of VirB3, which, in circular and cyclic sequence, facilitate serial insertion of individually recruited pilus subunits into the gap in the transverse VirB2 pentamer constituting the pilus base. A concurrent rise and rotation of the entire nascent pilus with each subunit incorporation event ensures that the incorporation cycles are geometrically consistent with each other and that the IMC/pilus symmetry mismatch is fully accommodated.

## RESULTS AND DISCUSSION

### Amber suppression allows for site-specific photo-crosslinking of intact R388 T4SS

We set out to substitute specific residues of the R388 T4SS with the non-canonical amino acid *p*-benzoyl-L-phenyl-alanine (Bpa) using amber suppression [Fig 1A]. According to published structures (Macé *et al*, 2022; Macé C Waksman, 2024), Bpa-substitution of the valine residue at position 252 (Val252) of the TrwI_B6_ subunits of the R388 T4SS leads to positioning of the resulting Bpa252 side chains within the inter-subunit interfaces of the TrwI_B6_ pentamer [Fig 1B]. Upon exposure to near-UV irradiation (365 nm wavelength), incorporated Bpa side chains become excited and capable of forming covalent bonds with proximal CH-groups [Fig 1C]. For the Val252Bpa-TrwI_B6_ T4SS variant, UV-irradiation is thus hypothesised to lead to formation of covalently linked TrwI_B6_ oligomers [Fig 1D] that are not separated into their constituent parts during SDS-PAGE analysis.

**Figure 1:**
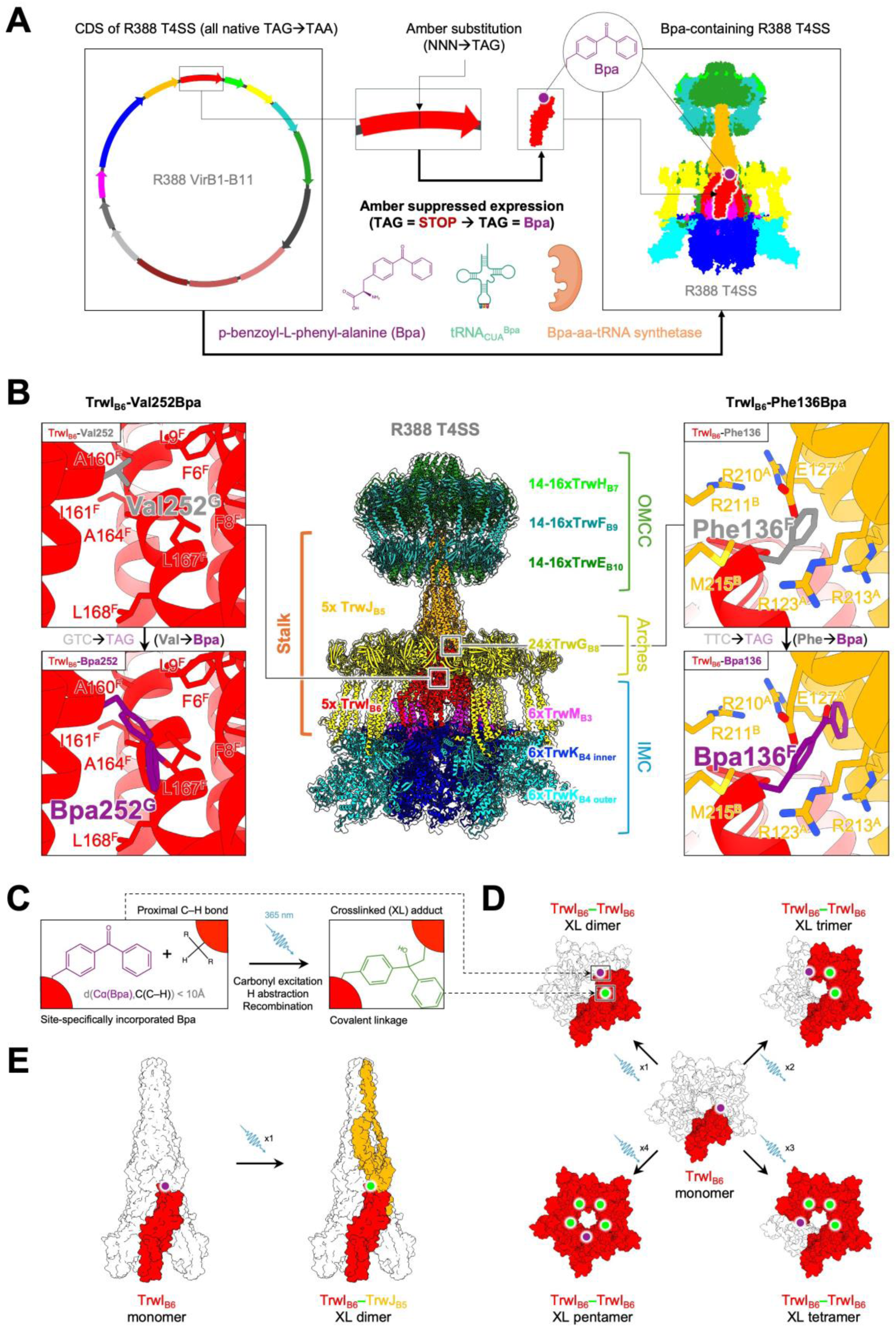
Photo-crosslinking of the R388 T4SS using site-specifically incorporated Bpa. **A)** Schematic drawing showing the strategy employed for producing intact R388 T4SS with site-specifically incorporated Bpa side chains. All natively occurring amber (TAG) codons of the R388 plasmid coding sequence (CDS) are modified to ochre (TAA) codons. Due to co-expression of the orthogonal aminoacyl-tRNA synthetase/suppressor tRNA (Bpa-aa-tRNA-synthetase/tRNA_CUA_^Bpa^) pair, amber-substitution (NNN → TAG) of specific codons in the R388 CDS leads to site-specific incorporation of Bpa supplied in the growth medium during expression. **B)** Overview of the composite model of the R388 T4SS in its resting, pre-activation state. The main architectural components (OMCC = Outer Membrane Core Complex; IMC = Inner Membrane Complex; Arches; Stalk) are shown. The identities of the different VirB protein components are indicated with colour coding. Two key positions of the Stalk segment are highlighted and shown in magnified view: Position 252 (left) and 136 (right) of two select subunit (G and F, respectively) within the TrwI_B6_ pentamer. Amber-substitution of the native codons leads to site-specific substitution of native side chains (grey sticks) for Bpa side chains (purple sticks). Only a single rotamer of the Bpa side chains are modelled for each position. **C)** Schematic drawing showing a simplified version of the Bpa side chain photo-crosslinking reaction (Dormán *et al*, 2016). Unreacted Bpa is shown in purple, while Bpa that has reacted with an incoming photon (shown in blue) and formed a covalent bond with the carbon atom of a nearby CH-group is shown in green. **D** and **E)** Expected patterns of crosslinked oligomer formation upon UV-treatment of the Val252Bpa- and Phe136Bpa-TrwI_B6_ R388 T4SS variants, respectively. Unreacted Bpa side chains are shown as purple dots, while Bpa side chains that have reacted with incoming photons (shown in blue) and formed a covalent bond with the carbon atom of a nearby CH-group are shown as green dots. The constituent parts of a selection of covalently crosslinked (XL) oligomers are highlighted, while the remaining elements of the overall architectural components that they form part of are outlined in grey.

Similarly, we also set out to substitute the phenylalanine residue at position 136 (Phe136) of the TrwI_B6_ subunits for Bpa. The resulting Bpa136 residues are positioned near the apex of the TrwI_B6_ pentamer, which in the published structures interfaces with the TrwJ_B5_ pentamer. Bpa substitution and subsequent UV-irradiation of the Phe136Bpa-TrwI_B6_ T4SS variant is therefore hypothesised to lead to formation of covalent links between the TrwI_B6_ and TrwJ_B5_ pentamers [Fig 1E]. Due to the multimeric nature of the R388 T4SS Stalk, even a single TrwI_B6_-to-TrwJ_B5_ crosslinking event will thus prevent the TrwI_B6_ and TrwJ_B5_ pentamers from separating.

To demonstrate the *in vitro* feasibility of this site-specific photo-crosslinking method, we recombinantly produced the Val252Bpa-TrwI_B6_ variant of the fully-assembled R388 T4SS by amber-suppressed expression [Fig 2A]. The complex was detergent-solubilised and intact T4SS purified by affinity chromatography [Fig 2B]. Aliquots of concentrated complex were then subjected to a time course of near-UV irradiation [Fig 2C]. At the beginning of the time course, only the C-terminally His-tagged TrwI_B6_ *mono*mer is detected. However, as UV irradiation was applied in increasing dose, *oligo*meric species appeared (expected electrophoretic mobilities indicated with cartoon representations), culminating in the formation of a species with electrophoretic mobility matching that expected for a TrwI_B6_ *penta*mer.

**Figure 2:**
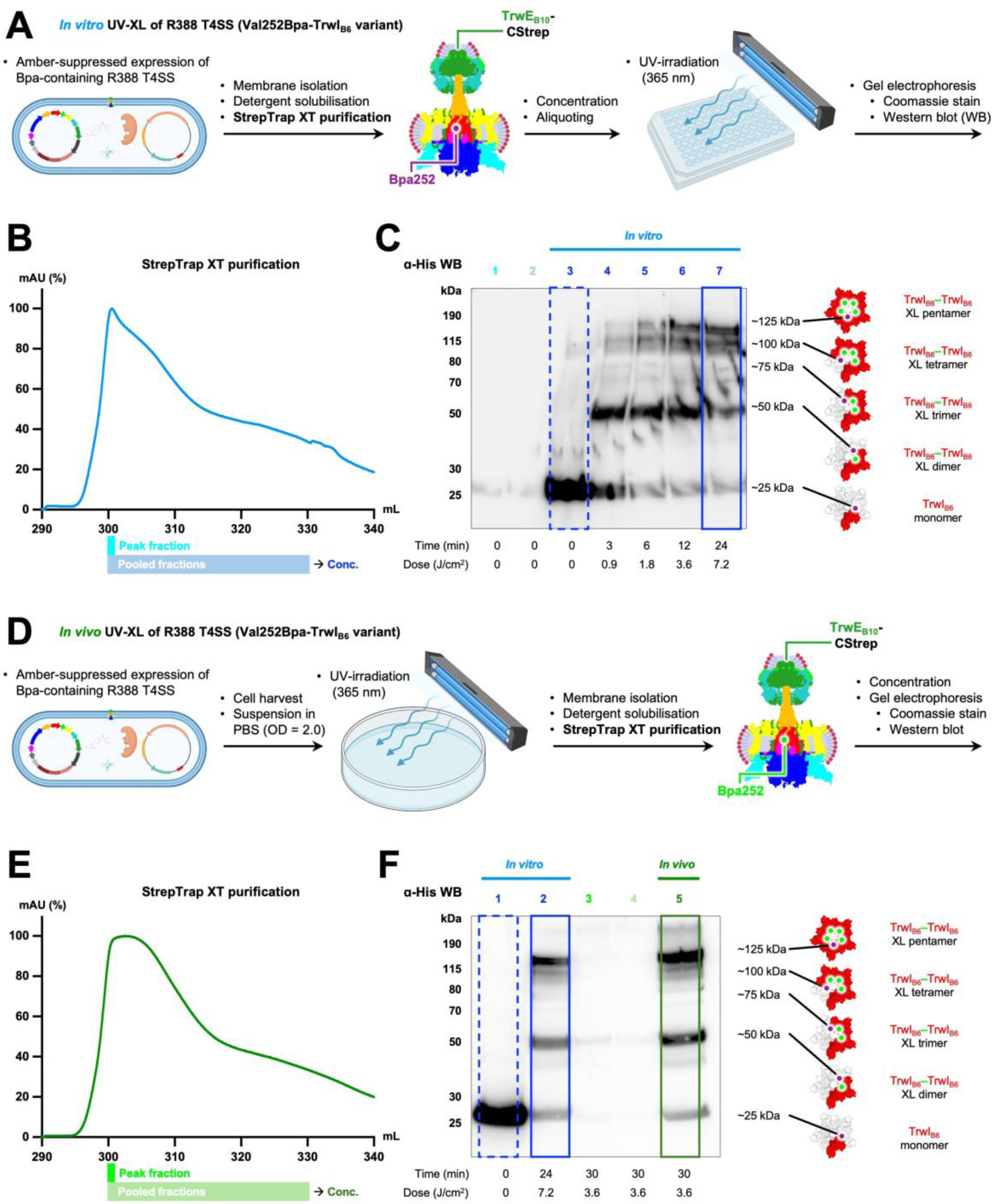
Photo-crosslinking of the Val252Bpa-TrwI_B6_ T4SS variant. **A)** Schematic diagram outlining the experimental procedure for *in vitro* UV-induced crosslinking (UV-XL). Complete complex is produced by amber-suppressed expression and intact, Bpa-containing R388 T4SS purified and concentrated before being subjected to UV-irradiation and electrophoretic analysis. **B)** StrepTrap XT affinity chromatogram from the purification of detergent-solubilised complex produced under amber-suppressing conditions. The peak fraction (Peak fraction) and the fractions pooled (Pooled fraction) and concentrated by tangential flow filtration to obtain a sufficiently concentrated, final sample (Conc.) are indicated. **C)** Anti-His Western blot analysis of the near-UV irradiation time course of purified, concentrated complex. The Bpa-containing component, TrwI_B6_, features a C-terminal His-tag, ensuring that only full-length protein is detected. Aliquots of ‘Peak fraction’ and ‘Pooled fractions’ (as shown in B)) are analysed in lanes 1 and 2, respectively. As is evident, extensive concentration (∼60-fold) of the StrepTrap XT eluate is instrumental for proper visualisation of TrwI_B6_. The hypothesised subunit compositions (and rough estimates of their corresponding electrophoretic mobilities) are shown for each of the covalently crosslinked oligomers expected to be formed upon UV-irradiation. **D)** Schematic diagram outlining the experimental procedure for *in vivo* photo-crosslinking. *Cells* producing complete R388 T4SS by amber-suppressed expression were subjected to UV-treatment *prior* to complex purification, concentration, and analysis by SDS-PAGE and Western blot. **E)** StrepTrap XT affinity chromatogram from the purification of *in vivo* UV-treated, detergent-solubilised complex produced under amber-suppressing conditions. The peak fraction and the fractions pooled and concentrated by tangential flow filtration are indicated. **F)** Side-by-side anti-His Western blot analysis of the aliquots of *in vitro* UV-treated, purified complex analysed in lanes 3 and 7 in panel C) (lanes 1 and 2, respectively), and aliquots of *in vivo* UV-treated, purified complex (lanes 3-5). As is evident, extensive concentration (∼60-fold) of the StrepTrap XT eluate is instrumental for proper visualisation of TrwI_B6_.

In parallel, the *in vivo* feasibility of this strategy was demonstrated by subjecting *cells* producing fully-assembled R388 T4SS by amber-suppressed expression to UV-irradiation *prior* to complex purification [Fig 2D]. As is evident from the affinity chromatogram [Fig 2E] and blots showing side-by-side analysis [Fig 2F], the results obtained from this *in vivo* experiment are practically indistinguishable from those obtained from the equivalent *in vitro* experiment (Fig 2F, compare lanes 2 and 5). In both the *in vivo* and *in vitro* experimental settings, we observe, as expected, substantial depletion of the TrwI_B6_ monomer, indicating efficient crosslinking.

Using mass spectrometry, we set out to determine the exact identities of the residues being crosslinked. Samples of untreated and UV-irradiated complex were processed and subjected to LC-MS/MS. From our spectral analyses, we were able to reliably identify precursor peptide ions with masses [Fig EV1A] and isotopic envelopes [Fig EV1B] matching those expected from *in silico* proteolysis and isotope simulations of the Bpa-containing complex.

It is expected that the proteolytic digests of UV-treated samples will contain a smaller proportion of Bpa-containing *mono*peptides than the proteolytic digests of the corresponding untreated samples. This relative depletion is due to the selective, chemical transformation of these *mono*peptides into *di*peptide adducts upon UV-induced crosslink formation. For crosslinking to be considered efficient, at least *one* monomer-to-monomer crosslinking event must occur per TrwI_B6_ pentamer. For a population of such pentameric assemblies, a more than 60% relative reduction of Bpa-containing monopeptide upon UV-treatment corresponds to an average of three independent crosslinking events having occurred within each pentamer. Assuming an underlying Poisson distribution with a mean population count of three (λ = 3) (Osborne *et al*, 2008), this corresponds to at least 95% of the pentameric assemblies containing at least one covalent crosslink. When we compared the mass spectra acquired from untreated and UV-irradiated samples, we indeed observed an 86.2% relative reduction of specific Bpa-containing *mono*peptides in the UV-treated samples, confirming efficient crosslinking [Fig EV2]. However, we have so far been unable to reliably detect any of the expected, Bpa-containing crosslink adducts directly. We hypothesise that low proteolytic susceptibility, high hydrophobicity, or other physicochemical properties of these dipeptide adducts effectively prevent their direct identification by LC-MS/MS (Kawashima *et al*, 2013).

### Covalent linkage of the two pentameric parts of the Stalk achieved by photo-crosslinking

We recombinantly produced the Phe136Bpa-TrwI_B6_ variant of fully-assembled R388 T4SS by amber-suppressed expression [Fig 3A]. The complex was detergent-solubilised and affinity-purified [Fig 3B], and aliquots of intact T4SS subjected to near-UV irradiation [Fig 3C]. This UV-treatment led to formation of at least two different oligomeric species (with apparent molecular weights of ∼45 and ∼65 kDa) that were not separated into their constituent parts during SDS-PAGE analysis [Fig 3C]. Western blotting confirmed that *both* bands contain *both* C-terminally Flag-tagged TrwJ_B5_ *and* C-terminally His-tagged TrwI_B6_.

**Figure 3:**
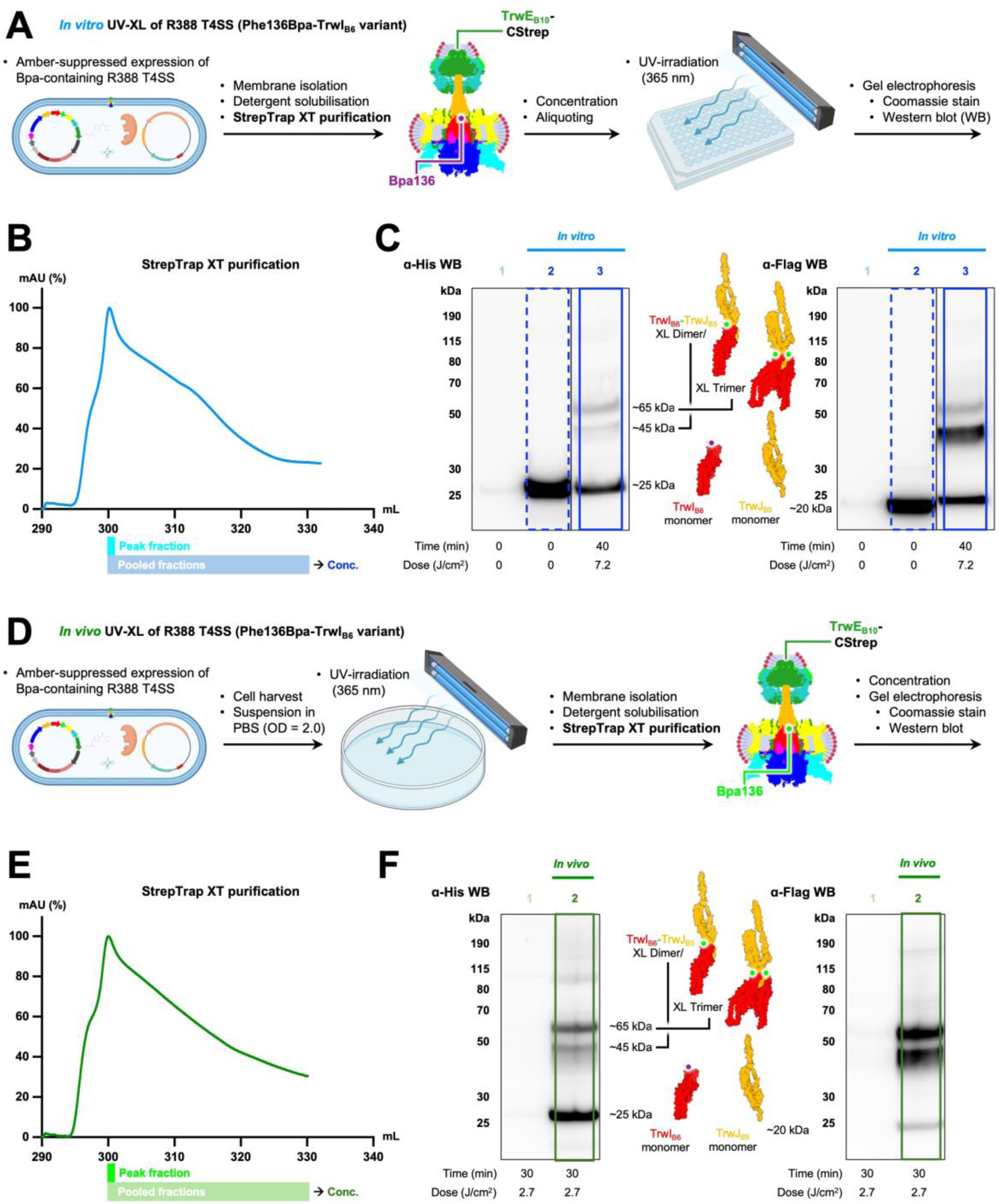
Photo-crosslinking of the Phe136Bpa-TrwI_B6_ variant. **A)** Schematic diagram outlining the experimental procedure for *in vitro* UV-induced crosslinking (UV-XL). Complete complex is produced by amber-suppressed expression and intact, Bpa-containing R388 T4SS purified and concentrated before being subjected to UV-irradiation and electrophoretic analysis. **B)** StrepTrap XT affinity chromatogram from the purification of detergent-solubilised complex produced under amber-suppressing conditions. The peak fraction (Peak fraction) and the fractions pooled (Pooled fractions) and concentrated by tangential flow filtration to obtain a sufficiently concentrated, final sample (Conc.) are indicated. **C)** Anti-His (left) and anti-Flag Western blot analysis of the near-UV irradiation treatment of purified, concentrated complex. The Bpa-containing component, TrwI_B6_, features a C-terminal His-tag, while TrwJ_B5_ features a C-terminal Flag-tag, ensuring that only full-length proteins are detected. An aliquot of ‘Pooled fractions’ (as shown in B)) is analysed in lane 1. As is evident, extensive concentration (∼60-fold) of the StrepTrap XT eluate is instrumental for proper visualisation of both His-tagged TrwI_B6_ and Flag-tagged TrwJ_B5_. The hypothesised subunit compositions (and rough estimates of their corresponding electrophoretic mobilities) are shown for each of the covalently crosslinked oligomers expected to be formed upon UV-irradiation. **D)** Schematic diagram outlining the experimental procedure for *in vivo* photo-crosslinking. *Cells* producing complete R388 T4SS by amber-suppressed expression were subjected to UV-treatment *prior* to complex purification, concentration, and analysis by SDS-PAGE and Western blot. **E)** StrepTrap XT affinity chromatogram from the purification of *in vivo* UV-treated, detergent-solubilised complex produced under amber-suppressing conditions. The peak fraction and the fractions pooled and concentrated by tangential flow filtration are indicated. **F)** Anti-His (left) and anti-Flag (right) Western blot analysis of aliquots of *in vivo* UV-treated, purified complex. An aliquot of ‘Pooled fractions’ (as shown in panel E)) is analysed in lane 1, while an aliquot of concentrated sample is analysed in lane 2. As is evident, extensive concentration (∼60-fold) of the StrepTrap XT eluate is instrumental for proper visualisation of both TrwI_B6_ and TrwJ_B5_.

In parallel, *cells* producing the Phe136Bpa-TrwI_B6_ variant of complete R388 T4SS by amber-suppressed expression were subjected to UV-irradiation *prior* to complex purification [Fig 3D]. The affinity chromatogram [Fig 3E] and Western blots [Fig 3F] of this *in vivo* experiment were similar to those obtained from the equivalent *in vitro* experiment. *In vivo* UV-treatment led to formation of the same two oligomeric species as in the *in vitro* experiment, and MS analysis [Fig EV3] confirmed that *both* bands contained *both* TrwJ_B5_ *and* TrwI_B6_.

LC-MS/MS analysis of gel bands corresponding to the crosslinked, oligomeric species yielded unambiguous evidence of site-specific Bpa-incorporation at both precursor level [Fig 4A-B] and fragmentation level [Fig 4C]. In addition, we observe a 66.4% relative reduction of Bpa-containing monopeptides upon UV-treatment [Fig EV2] and thus confirm efficient crosslink formation (see above). However, our analysis did not yield reliable, *direct* identification of crosslinked, *di*peptide adducts. Given their respective electrophoretic mobilities, the smaller oligomeric species (∼45 kDa) thus appears to correspond to a covalently crosslinked, chimeric TrwJ_B5_-TrwI_B6_ dimer, while the larger species (∼65 kDa) likely corresponds to a chimeric, covalently linked *tri*mer featuring an additional copy of TrwI_B6_.

**Figure 4:**
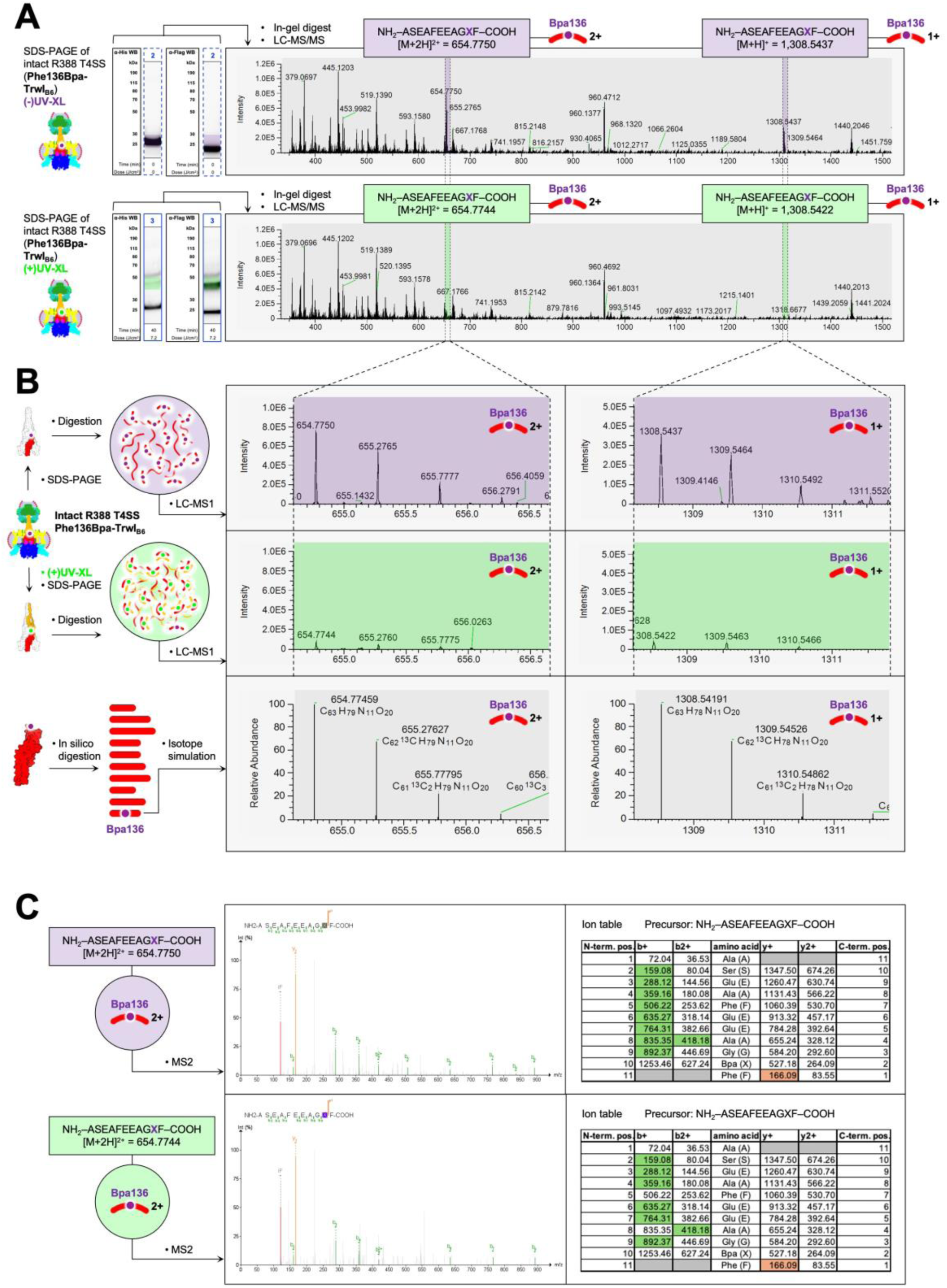
LC-MS/MS analysis confirming site-specific Bpa-incorporation and photo-crosslinking of the R388 T4SS Phe13cBpa-TrwI_B6_ variant. **A)** Representative precursor (MS1) spectra obtained from LC-MS/MS analyses of in-gel digestions of dissected gel bands from SDS-PAGE analysis of specific samples of untreated (purple) and UV-treated (green) complex shown in Fig 3C. Based on *in silico* trypsin/chymotrypsin digestion of TrwI_B6_, we determined the exact masses of the singly ([M+H]^+^) and doubly ([M+2H]^2+^) charged species of the proteolytic peptide NH_2_-ASEAFEEAGXF-COOH (spanning sequence positions 127-137 and featuring the expected Phe (F) to Bpa (X) substitution at position 136). Automated peak detection identified peaks corresponding to these masses with retention times of the corresponding ion chromatograms being similar (1,430.64 s and 1,427.85 s, respectively) for the untreated and UV-treated samples (see Fig EV2). **B)** Magnified views of selected peaks of the mass spectra shown in panel A). The observed isotopic envelopes fit with the simulated isotopic envelope of the singly ([M+H]^+^) and doubly ([M+2H]^2+^) charged species of the NH_2_-ASEAFEEAGXF -COOH peptide. Selective, relative depletion (quantified in Fig EV2) of this Bpa-containing peptide in the UV-treated sample (green) relative to the untreated sample (purple) is evident. **C)** Representative fragmentation (MS2) spectra and corresponding ion tables obtained from LC-MS/MS analysis of the indicated samples analysed above. Fragmentation spectra and corresponding ion tables are only shown for the doubly charged species.

We thus show that both *in vitro* and *in vivo* UV-treatment of the Phe136Bpa-TrwI_B6_ T4SS variant results in robust formation of expected, covalently crosslinked, chimeric TrwJ_B5_-TrwI_B6_ species. Intriguingly, we demonstrate that UV-irradiation of *one* variant of the R388 T4SS, with Bpa incorporated at a *single* position, can give rise to multiple different crosslinked adducts. Since the experiments are conducted on *intact* complex (including intact complex in an *in vivo* setting), our results likely reflect *bona fide*, biologically relevant, dynamic rearrangement of the Bpa-containing subunits.

### Crosslinking TrwI_B6_ to TrwJ_B5_ has only negligible effects on conjugation efficiency

Since pilus elaboration is essential for conjugative transfer by the R388 T4SS (Vadakkepat *et al*, 2024), any obstruction of pilus biogenesis will result in a decrease in conjugative transfer efficiency. According to prevailing models of pilus biogenesis (Macé *et al*, 2022; Macé C Waksman, 2024; Costa *et al*, 2023; Paillard *et al*, 2025), the TrwJ_B5_ and TrwI_B6_ assemblies of the R388 T4SS Stalk become completely separated during pilus growth: The TrwI_B6_ pentamer remains anchored to the inner membrane, while the TrwJ_B5_ pentamer becomes positioned at the tip of the growing pilus. Consequently, covalently linking the two pentameric parts of the Stalk and thus preventing their separation should therefore have a proportionally strong, negative effect on conjugative transfer efficiency.

We set out to investigate the functional effects of UV-induced crosslinking of our Bpa-containing R388 T4SS variants using conjugation assays (Redzej *et al*, 2017; Macé *et al*, 2022) [Fig 5A]. We constructed a three-component plasmid system for the purpose of investigating the effects of Bpa-incorporation and UV-induced crosslinking on *in vivo* conjugative plasmid transfer by the R388 T4SS (see Materials and Methods for details). For the conjugation assays, donor cells encoding either an amber-substituted variant or the amber-less control variant of complete R388 T4SS were cultivated under identical, amber-suppressing conditions. Duplicate aliquots of these donor cell strains were then either left untreated or subjected to UV-irradiation under conditions demonstrated (in this study) to induce robust formation of expected crosslinks. Since the R388 T4SS only effectuates conjugation under solid-support conditions (Bradley, 1980; Bolland *et al*, 1990), the donor cell aliquots were then assayed for their capacity to perform conjugative transfer of a donor plasmid to recipient cells by solid-support mating.

**Figure 5:**
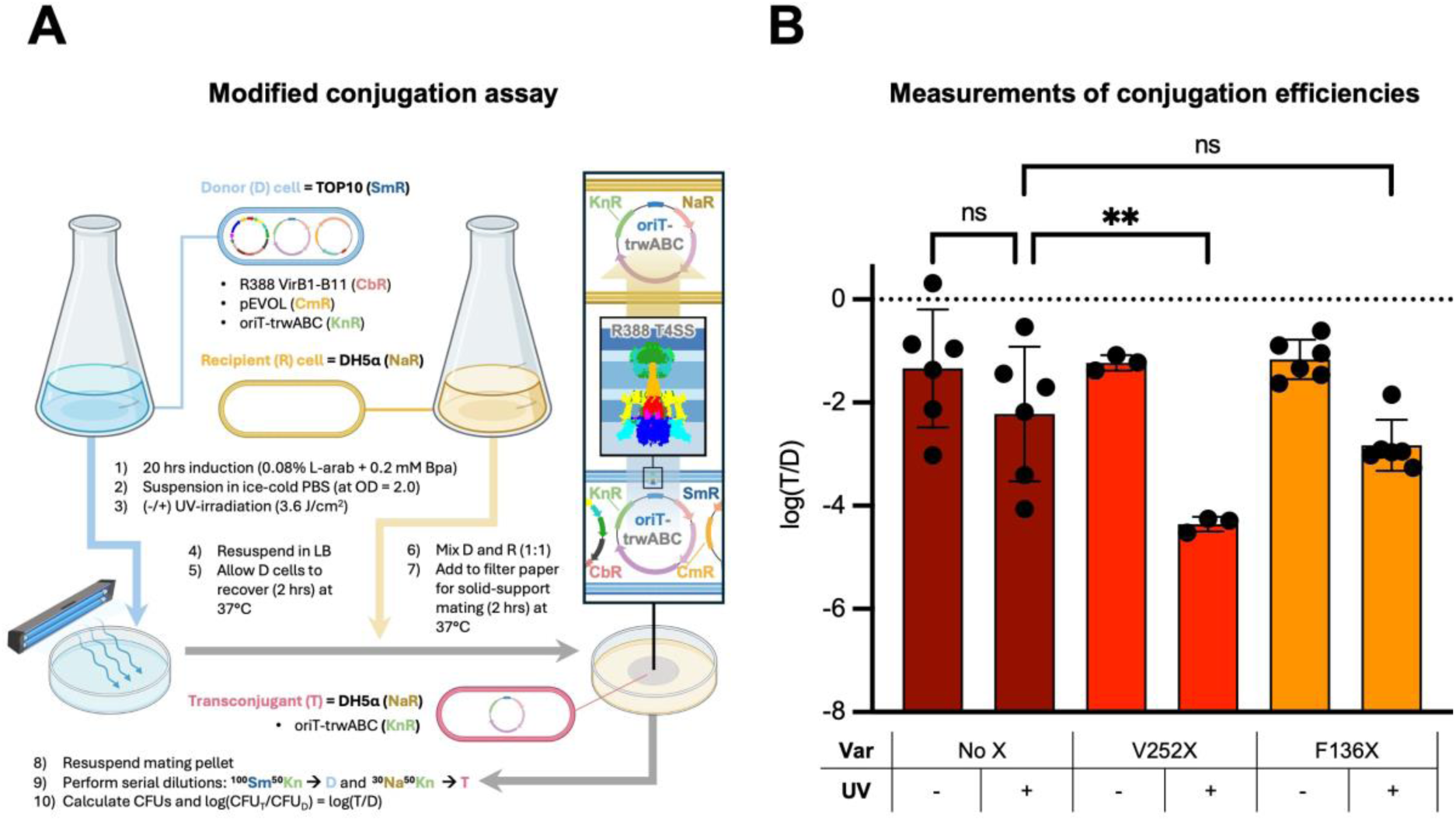
Investigating the effects of UV-irradiation on the conjugation efficiency of donor cells expressing different variants of complete R388 T4SS. **A)** Schematic diagram outlining the experimental procedure for the modified conjugation assay (see Materials and Methods for details). The insert (outlined in black) shows a schematic drawing of the conjugative process leading to formation of transconjugants (T) during solid-support mating of donor (D) and recipient (R) cells. **B)** Graph displaying the results of the modified conjugation assays. Mean conjugation efficiencies, calculated as log(T/D) (see Materials and Methods for details) of untreated (-) and UV-treated (+) aliquots of the indicated donor cell variants (Var), are shown as coloured bars. Results of individual, independent experiments (n ≥ 3) are shown as black dots and standard deviations of means indicated with error bars. The y-axis was arbitrarily truncated at -8 (corresponding to a conjugation efficiency of 10^-8^ T/D), which reflects the detection limit of the assay. The brackets above the bars report the results of the chosen Bonferroni’s multiple comparisons test (see Materials and Methods for details) comparing the indicated pairs of means using a standard alpha threshold of 0.05 (95% confidence interval) and correction for multiplicity. Asterisks indicate statistical significance, while ’ns’ indicates ’no significance’. Multiplicity-adjusted P-values: ’No X (+)’ vs. ’No X (-)’, P = 0.2464 (ns); ’V252X (+)’ vs. ’No X (+)’, P = 0.0046 (**); ’F136X (+)’ vs. ’No X (+)’, P = 0.6821 (ns).

All assayed strains exhibited similar expression levels of complete T4SS [Fig EV4], which allowed us to directly compare the relative effect of UV-irradiation (and UV-induced crosslinking) on the conjugation efficiency of the different variants.

Previous studies have shown that near-UV illumination affects bacterial physiology by shutting down, among other cellular processes, protein synthesis (Waksman *et al*, 1984). The same studies have shown that the effect is reversible, with bacteria displaying recovery of cellular activities after 2 hours. We therefore opted for a modified conjugation assay protocol, in which donor cells are allowed to recover for 2 hours after the time of UV-treatment. In the bar graph displaying the conjugation assay results [Fig 5B], (+) therefore denotes ’UV-treatment followed by a 2-hour recovery period’, while (-) denotes ’no UV-treatment’ before measurement of the conjugation efficiencies of aliquots of cells expressing the indicated variants (Var) of the R388 T4SS.

From our results, we note first that Bpa-incorporation *itself* does not affect conjugation efficiency, as the amber-substituted variants (’V252X’ and ’F136X’, expressing Val252Bpa-TrwI_B6_ and Phe136Bpa-TrwI_B6_, respectively) exhibit the same absolute, mean conjugation rates as the amber-less (’No X’) control variant (Fig 5B, compare ’V252X (-)’ and ’F136X (-) to ’No X (-)’). We note next a small reduction in conjugation efficiency upon UV-irradiation of the amber-less control variant (Fig 5B, compare ’No X (+)’ and ’No X (-)’). Given that no Bpa-incorporation, and thus no UV-induced crosslinking, can be expected to occur for this amber-less control variant, this reduction therefore represents the background effect of UV-irradiation *in itself* on conjugation – even in the absence of any Bpa-crosslinking. We therefore conclude that, in order to properly correct for this background effect, the effects of UV-irradiation on the investigated variants must be assessed by comparing the resulting conjugation rates (’V252X (+)’ and ’F136X (+)’) to this background-adjusted control rate (’No X (+)’).

When compared to the control strain, we observe a strong, negative effect of UV-irradiation on cells expressing the ’V252X’ variant (a 136-fold reduction in conjugation efficiency when comparing ’V252X (+)’ to ’No X (+)’ in Fig 5B). UV-irradiation of this variant has been demonstrated to covalently link the TrwI_B6_ subunits of the T4SS Stalk, and our results thus show that this inter-subunit crosslinking of the TrwI_B6_ pentamer interferes strongly with conjugative plasmid transfer. A Bonferroni’s multiple comparisons test indeed confirms that there is a statistically significant difference (**; multiplicity-adjusted P = 0.0046) between the mean conjugation rates of ’V252X (+)’ and ’No X (+)’, meaning that there is a significant, negative effect of UV-induced inter-TrwI_B6_ crosslinking on the conjugation efficiency of the ’V252X’ variant [Fig 5B]. This observation indicates that relative subunit rearrangement within the TrwI_B6_ pentamer is required for conjugation and indicates that the TrwI_B6_ pentamer does not simply serve as a *passive* platform for pilus assembly, but rather is dynamically rearranged during the conjugation process.

To our surprise, we observed only a relatively minor effect of UV-irradiation on the conjugation efficiency of cells expressing the ’F136X’ variant (a 4-fold reduction in conjugation efficiency when comparing ’F136X (+)’ to ’No X (+)’ in Fig 5B). We have demonstrated that UV-irradiation of this variant leads to covalent crosslinking of the two pentameric TrwI_B6_ and TrwJ_B5_ Stalk parts, which would prevent their separation during pilus growth. We therefore expected that UV-irradiation would severely limit conjugative donor plasmid transfer by completely inhibiting pilus biogenesis. However, our statistical analysis (see above) shows that there is no statistically significant difference (ns; multiplicity-adjusted P = 0.6821) between the mean conjugation rates of ’F136X (+)’ and ’No X (+)’, meaning that there is no significant effect of UV-induced TrwI_B6_-TrwJ_B5_ crosslinking on the conjugation efficiency of the ’F136X’ variant [Fig 5B]. Our results thus show that UV-induced, covalent crosslinking of the two Stalk pentamers, which prevents their separation, does *not* inhibit conjugation, and permits regular levels of pilus formation and conjugative transfer. This observation implies that the TrwJ_B5_ and TrwI_B6_ pentamers *remain* closely associated with each other during pilus biogenesis, and, since VirB5 has been shown to locate at the pilus tip (Aly C Baron, 2007), implies that TrwI_B6_ is *also* a tip protein.

### Experimental evidence of VirB2 recruitment and binding to VirB3 of the IMC

Our previously proposed pilus biogenesis model (Macé *et al*, 2022) assumed that TrwI_B6_ serves as a membrane-anchored base onto which the pilus is built. However, our new results contradict this model, and, therefore, a new assembly base needs to be proposed. In a previous study (Macé *et al*, 2022), we showed that another candidate for TrwL_B2_-recruitment other than TrwI_B6_ was TrwM_B3_. Direct interaction between TrwL_B2_ and TrwM_B3_ was initially inferred from co-evolution analyses (Macé *et al*, 2022) carried out using TrRosetta (Song *et al*, 2013). However, direct experimental evidence that VirB3 was directly involved in pilus biogenesis was lacking. We have previously hypothesised (Macé C Waksman, 2024) that a loop-shaped density observed close to TrwM_B3_ in the cryo-EM map of the R388 IMC protomer (EMD-19485) might correspond to TrwL_B2_, which exhibits a distinct helix-loop-helix hairpin morphology (Vadakkepat *et al*, 2024). However, the density for side chains was ill-defined, and, therefore, unambiguous identification of the protein contributing to this density was impossible.

The R388 and pKM101 conjugative plasmids and their respective T4SSs are close homologues (Bolland *et al*, 1990; Paterson C Iyer, 1997; Paterson *et al*, 1999), with many of their components being functionally interchangeable (Bolland *et al*, 1990; Llosa *et al*, 2003; Gordon *et al*, 2017). Spurred on by the close homology between the two systems, we applied cryo-EM to samples of the T4SS encoded by the pKM101 plasmid to solve the pKM101 IMC protomer structure in the hope that the density for the VirB2 homologue of that system would be more clearly interpretable. Samples of the pKM101 T4SS were produced and prepared like the samples used for structure determination of the R388 T4SS in its resting, pre-activation state (Macé *et al*, 2022). Our single particle reconstruction of the pKM101 IMC protomer [Fig EV5] revealed a loop-shaped density similarly positioned to the one observed in the structure of the R388 T4SS, but of improved quality and interpretability [Fig EV6A]. Distinct density corresponding to bulky side chains and proline-induced kinks allowed for unequivocal model building of residues 45-89 of pKM101 TraM_B2_, corresponding to two thirds of the mature pilin protein (residues 28-97). Based on this model building, the equivalent residues (62-106) of the pilin moiety of the model of the R388 TrwL_B2_-TrwM_B3_ complex obtained using AlphaFold 3 (Abramson *et al*, 2024) (AF3; see below) was fitted into the published map of the R388 IMC protomer (Macé C Waksman, 2024), which yielded good correspondence regarding shape and location of bulky side chains [Fig EV6B].

In our experimentally determined structure of the pKM101 IMC protomer [Fig 6A], TraM_B2_ can be seen to clearly interact directly with TraA_B3_ [Fig 6B and EV6A]. The interface between TraM_B2_ and TraA_B3_ is bipartite and involving residues for which distinct side chain density is observed, with 12.9% and 4.8% of the solvent accessible area of the respective components being buried.

**Figure 6:**
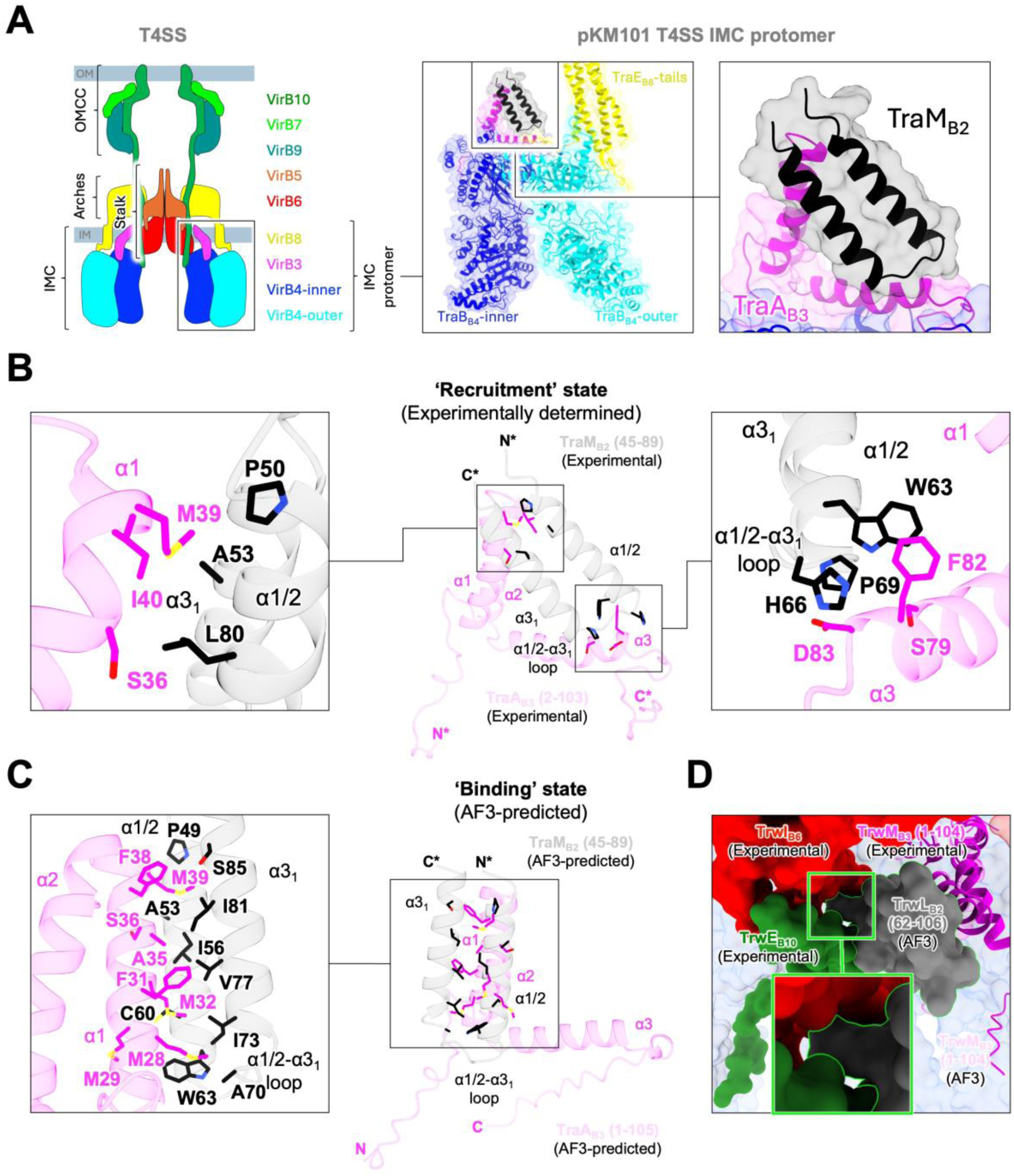
Structural characterisation of the pKM101 T4SS IMC protomer using cryo-EM and AF3. **A)** The experimentally determined structure of the pKM101 T4SS IMC protomer. Left: Schematic representation of a generic T4SS in its resting, pre-activation state. The main architectural components (OMCC = Outer Membrane Core Complex; IMC = Inner Membrane Complex; Arches; Stalk) are shown. The identities of the different VirB protein components are indicated with colour coding. Middle: Highlighted view of the colour coded pKM101 T4SS IMC protomer model (cartoon and surface representation). Right: Highlighted view of the IMC protomer model (surface representation) showing the built models of the pilin protein, TraM_B2_ (black cartoon), and TraA_B3_ (magenta cartoon). **B)** Visualisation and characterisation of the bipartite interaction interface between pKM101 TraM_B2_ (black) and TraA_B3_ (magenta) in the experimentally determined (Experimental) ‘Recruitment state’ configuration. The buried surface area was calculated using PDBePISA (available at https://www.ebi.ac.uk/pdbe/pisa/) (12.9% for TraM_B2_; 4.8% for TraA_B3_) and side chains determined to be directly involved in interfacial interactions are shown as sticks. **C)** Visualisation and characterisation of the unipartite interaction interface between pKM101 TraM_B2_ (black) and TraA_B3_ (magenta) in the AF3-predicted (AF3-predicted) ‘Binding state’ configuration. The buried surface area was calculated using PDBePISA (15.8% for TraM_B2_; 6.6% for TraA_B3_) and side chains determined to be directly involved in interfacial interactions are shown as sticks. **D)** Highlighted view of the R388 IMC+Stalk model. The AF3-predicted (AF3) structure of the TrwL_B2_-TrwM_B3_ binary complex (respectively, grey surface outlined in green and faded magenta cartoon) has been aligned along a select TrwM_B3_ component (magenta cartoon) of the R388 IMC+Stalk model. The nearest TrwI_B6_ (red surface) and TrwE_B10_ (green surface) components are highlighted. The insert (box with green outline) shows a magnified view of the steric clash.

### AF3-modelling of the VirB3 interaction with VirB2 reveals a binding site on VirB3 different from the one observed experimentally

AF3 can be used to predict interactions between proteins with great accuracy and reliability. We used AF3 to obtain a model (AF3-predicted) of the interaction between pKM101 TraM_B2_ and TraA_B3_, and compared it to our experimentally determined model (Experimental) [Fig EV6C]. For the AF3-predicted model, we used the full, mature pilin protein (residues 28-97), which we then subsequently, for the sake of direct comparison with the Experimental model, manually truncated to match the sequence stretches represented by density in the IMC protomer cryo-EM map (residues 45-89). The residues found to be directly involved in the Experimental TraM_B2_-TraA_B3_ interaction [Fig 6B] were confidently modelled by AF3 [Fig EV6C]. The AF3-predicted interface between TraM_B2_ and TraA_B3_ is *uni*partite [Fig 6C], with 15.8% and 6.6% of the solvent accessible area of the respective components being buried. The Experimental [Fig 6A-B and EV6A] and AF3-predicted [Fig 6C and EV6C] configurations of the pKM101 TraM_B2_-TraA_B3_ interaction are related to each other by a ∼40° pivot of the TraM_B2_ moiety. The same is the case for the equivalent Experimental and AF3-predicted configurations of the R388 TrwL_B2_-TrwM_B3_ interaction [Fig EV6D]. These two binding configurations appear to be mutually exclusive, indicating that they correspond to distinct, sequential steps in pilus subunit recruitment and binding (see below).

Intriguingly, aligning the AF3-predicted structure of the R388 TrwL_B2_-TrwM_B3_ binary complex along the modelled TrwM_B3_ components of the R388 Stalk+IMC cryo-EM map (EMD-19488) leads to a steric clash between TrwL_B2_ and the transmembrane helices of TrwI_B6_ and TrwE_B10_ [Fig 6D]. This suggests that TrwI_B6_ and TrwE_B10_, in the resting, pre-activation state of the R388 T4SS, are *sterically obstructing* a potential transition from the Experimental TrwL_B2_-TrwM_B3_ configuration, representing a ‘Recruitment state’ of TrwL_B2_ on TrwM_B3_, to the AF3-predicted configuration, which may represent a *bona fide* ‘Binding state’ of TrwL_B2_ on TrwM_B3_.

Combined with the results of our crosslinking experiments, which imply that TrwI_B6_ co-locates with TrwJ_B5_ at the pilus tip, this indicates that TrwI_B6_ is likely, immediately prior to pilus biogenesis, rearranged. This rearrangement allows for the transition of TrwL_B2_ from a ‘Recruitment’ to a ‘Binding’ interaction with the TrwM_B3_ components of the IMC, which therefore appear to serve as the base for assembly of the conjugative pilus. Owing to the strong homology between the two systems [Fig EV6E], an equivalent mechanism is likely to be at play in the pKM101 T4SS.

### A revised model of conjugative pilus biogenesis

Our findings are incompatible with the prevailing model of conjugative pilus biogenesis (Macé *et al*, 2022; Costa *et al*, 2023), in which the VirB5 and VirB6 pentamers of the T4SS Stalk must separate during pilus assembly in order for VirB6 to act as a platform for pilus assembly, while VirB5 is projected at the distal pilus end [Fig 7A]. Covalently linking the two pentamers has only a modest effect on conjugation, which indicates that separation of the VirB5 and VirB6 pentamers is in fact *not* required for pilus biogenesis and conjugative transfer to occur. We note that our findings are consistent with the recently deposited cryo-EM structure of the R388 conjugative pilus tip complex that shows TrwI_B6_ “sandwiched” between TrwJ_B5_ and the helical TrwL_B2_ polymer (preprint: Paillard *et al*, 2026) . While our crosslinking results provide functional validation for this structure, the latter provides structural validation for our results.

**Figure 7:**
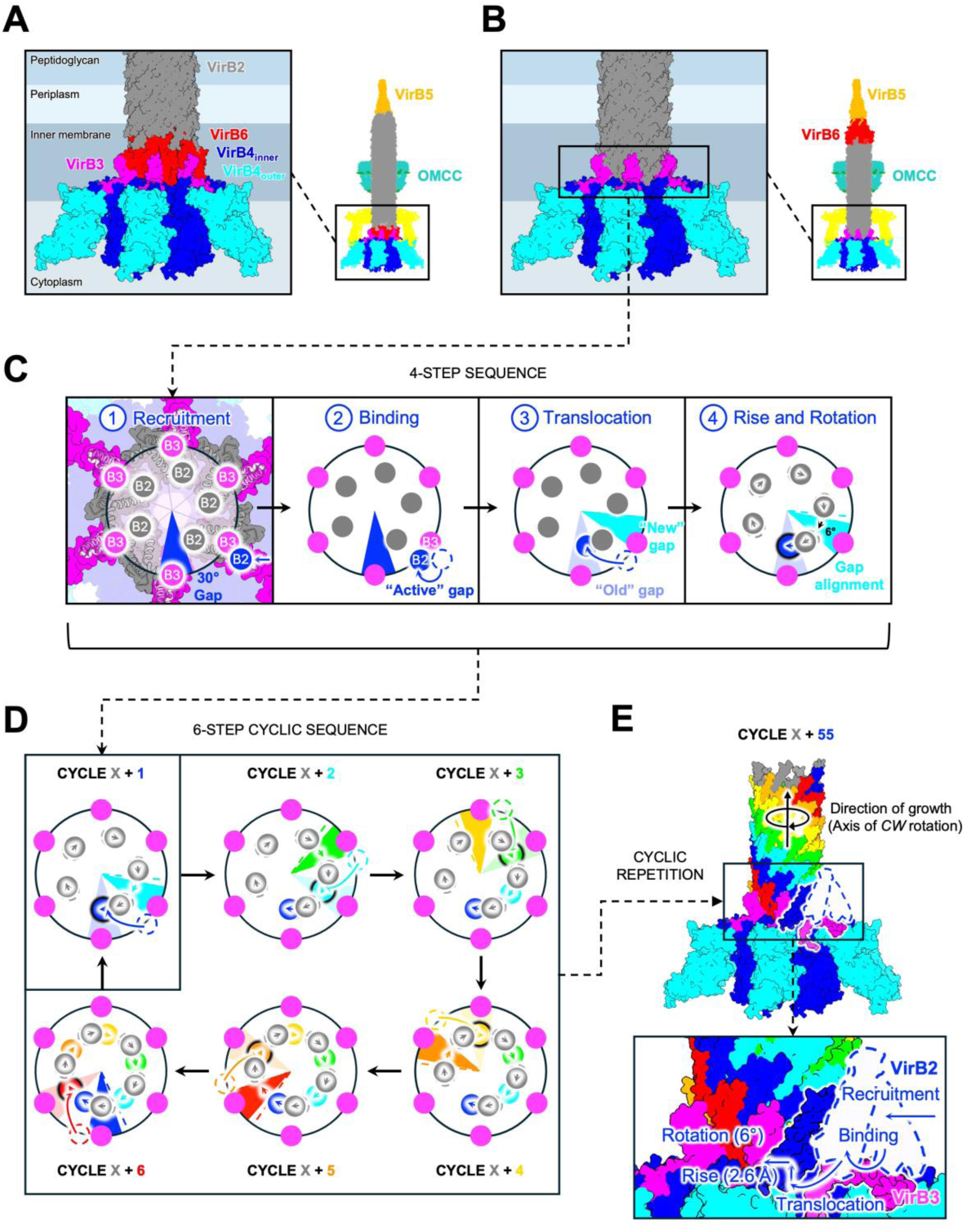
Revised model of conjugative pilus biogenesis. **A** and **B)** Composite models of the R388 conjugative pilus (PDB: 8S6H, pilin protein (TrwL_B2_) shown in grey) and components of the hexameric R388 IMC model (PDB: 7O41, TrwM_B3_ shown in magenta, TrwK_B4_ shown in shades of blue) representing the prevailing (left) and revised (right) models of conjugative pilus biogenesis, respectively. All docking was done manually based on size, shape, and symmetry complementarity between the different components (see Materials and Methods for details). The Arches and the OMCC (PDB: 7O3J/7O3T, TrwG_B8_ shown in yellow, TrwH_B7_, TrwF_B9_, TrwE_B10_ shown in shades of green) have been sliced vertically in the full views and completely removed from the magnified views for the sake of clarity. Rough estimates of the relative positions of the different segments of the donor cell envelope are shown in shades of blue in the magnified views. **C)** Schematic outline of the four steps of the cyclic pilus subunit incorporation sequence (incorporation cycle). The nascent pilus is viewed from the extracellular side along the longitudinal axis, with the pilus growing towards the viewer. Surface and cartoon representations of the six IMC copies of TrwM_B3_ (magenta) are highlighted, and their mean geometric positions (along the periphery of an imagined circle (black), with adjacent TrwM_B3_ copies being related by a 60° turn around the centre) are marked with labelled dots. Likewise, surface and cartoon representations of the five (last to be incorporated) subunits of the bottom-most pentamer of the pilin protein, TrwL_B2_ (grey), are shown, and their mean geometric positions marked with labelled dots. The relevant 30° gaps (i.e. incorporation sites) are shown as coloured wedges. The “Old” (blue) and “New” (cyan) gaps are related to each other by a 66° turn, reflecting the relation between the adjacent subunits of the pentamer. Gap alignment and movements of the pilus subunits are indicated with coloured arrows (signifying a 6° turn). The IMC (represented by the six copies of TrwM_B3_) is kept as a constant frame of reference. Note that the blue pilus subunit being incorporated in the shown incorporation cycle (black glow upon incorporation) is inserted *below* the grey pilus subunits, and that this incorporation involves the entire nascent pilus rotating by 6° (*CW* for the viewer) and rising 2.6 Å (towards the viewer). **D)** Schematic outline of six sequential steps (each containing the four steps shown in panel C)) of the cyclic sequence leading to pilus elaboration. View and colour scheme as in panel C). For sake of clarity, only steps 3-4 in panel C) have been drawn. **E)** Composite structural model (surface representation) of relevant IMC components and the conjugative pilus at the end of incorporation cycle X + 55. Colour scheme as in panel D). Magnified side view depicts the four steps shown in panel C). The copy of TrwL_B2_ being incorporated and the copy of TrwM_B3_ actively involved in this incorporation are highlighted and moved to the foreground for the sake of clarity. The relative movements are shown with blue arrows. The ‘Recruitment’ and ‘Binding’ configurations of the TrwL_B2_-TrwM_B3_ interaction (see Materials and Methods for details) are shown as white shapes outlined with a dashed, blue line.

Considering this new evidence, we set out to model an alternative arrangement of the R388 pilus relative to the R388 T4SS. Guided by the distinct size and shape complementarity between the convex end of the pilus structure and the concave architecture of the periplasm-facing side of the R388 T4SS IMC, we manually docked the fivefold-symmetrical pilus structure directly onto the sixfold-symmetrical structure of the IMC [Fig 7B]. This modelling left us with the seemingly intractable problem of a distinct sixfold-to-fivefold symmetry mismatch. However, careful examination of the R388 pilus structure revealed that the fivefold symmetry of the pilus is effectively already partly transitioned towards sixfold symmetry: Indeed, the relative arrangement of the longitudinal strands causes the subunits of the resulting transverse pentamers to be related to each other by a ∼66° twist and ∼2.6 Å rise (constituting a 1-start helical symmetry operation applied along the entire pilus), which introduces a ∼30° gap with a ∼13 Å rise in each pilus subunit pentamer (Vadakkepat *et al*, 2024) [Fig 7C].

In the structure of the R388 T4SS transfer apparatus (Macé *et al*, 2022), six copies of the IMC protein TrwM_B3_ are arranged in a ring. In our model, each of the six copies of TrwM_B3_ serves as a pilus subunit incorporation station: In a multistep, cyclic sequence, individual pilus subunits are recruited to the surface of TrwM_B3_ (as per modelling described above) and, ultimately, inserted into the gap of the bottom-most transverse pentamer of the nascent pilus. Each of these incorporation cycles culminates in a discrete rise and rotation of the *entire* nascent pilus, which ensures proper alignment of the newly repositioned gap with the adjacent TrwM_B3_ incorporation station, and provides distinct directionality to the pilus growth. In a revolving manner, incorporation events thus effectively propagate around the six-membered ring of the TrwM_B3_ incorporation stations [Fig 7D], leading to gradual, geometrically consistent assembly and elaboration of the conjugative pilus [Fig 7E and Mov S1].

Crucially, the 6° rotation and 2.6 Å rise per incorporation cycle resolves the mismatch between the 60° turn relating the individual TrwM_B3_ recruitment stations and the 66° twist and 2.6 Å rise relating the individual subunits of the transverse TrwL_B2_ pentamer [Fig EV7]. Our model thus explains how a central symmetry mismatch between the hexameric IMC and the pentameric pilus is not just accommodated, but in fact mechanistically utilised during conjugative pilus biogenesis.

## CONCLUSIONS

In this study, we demonstrate the feasibility of using site-specific Bpa-incorporation and subsequent UV-induced crosslink formation to investigate the structural dynamics of the R388 T4SS. We demonstrate that Bpa residues can be reliably introduced at key positions within this complex nanomachinery without affecting its structure and function. We show that subsequent UV-treatment of the intact complex results in robust formation of expected, oligomeric, covalently crosslinked species.

Inter-subunit crosslinking of the TrwI_B6_ pentamer of intact R388 T4SS is shown to strongly impede conjugative transfer, indicating that this pentameric Stalk component is dynamically rearranged during the conjugation process. Surprisingly, covalently linking the TrwI_B6_ pentamer to the other half of the Stalk, a pentamer of TrwJ_B5_, is shown to have only a minor effect on the conjugation efficiency of the R388 T4SS. This observation conflicts with prevailing models of conjugative pilus biogenesis and indicates that the two pentameric components of the Stalk do not separate during pilus biogenesis.

Cryo-EM reveals direct interactions between the pilin protein, TraM_B2_, and TraA_B3_ of the pKM101 T4SS IMC, which sheds light on the identity of previously uninterpretable density in the published cryo-EM maps of the R388 T4SS, which can now confidently be attributed to TrwL_B2_. Combined with AF3-modelling, our results indicate that, in both the R388 and pKM101 secretion systems, the VirB3 components of the IMC form a recruitment base for assembly of the conjugative pilus, which remains sterically obstructed by VirB6 and VirB10 in the resting, pre-activation state.

In the light of this new evidence, we propose a revised model of conjugative pilus biogenesis, which addresses how a central, sixfold-to-fivefold symmetry mismatch between the T4SS IMC and the conjugative pilus is accommodated and even mechanistically utilised. In our model, the six IMC copies of VirB3 function as individual incorporation stations that interact directly with and coordinate the serial insertion of incoming VirB2 subunits at the pilus base. Central to the proposed mechanism is the discrete rotation and rise of the entire nascent pilus accompanying each pilus subunit incorporation event. The resulting, screwlike motion ensures that the incorporation cycles are geometrically consistent with each other and provides unequivocal directionality to the pilus growth.

Symmetry mismatches such as the ones observed in T4SSs (Macé C Waksman, 2024) are ubiquitous in macromolecular complexes, where they provide necessary flexibility required for normal function (Zehra *et al*, 2023). Direct sixfold-to-fivefold symmetry mismatches have been observed in other secretion systems (Nazarov *et al*, 2018; Rapisarda *et al*, 2019), with some even involving transitions from rotational to helical symmetry (Tan *et al*, 2021).

Apart from symmetry *mismatches*, transient symmetry *disruptions* during conformational cycling are also widely observed among macromolecular machines (Bohmer *et al*, 2023). In complexes featuring hexameric ATPase assemblies like those found in the R388 T4SS, a rotary mechanism involving cycling through various asymmetrical states is widely employed for translocation of DNA substrates (Lyubimov *et al*, 2012; Sun *et al*, 2017; Jean *et al*, 2020). Similar ATPase-driven rearrangements have been proposed as the underlying mechanistic framework of the assembly of Type IV pili (Nayak *et al*, 2022; Yen *et al*, 2025; Guo *et al*, 2026). In particular, the suggested molecular mechanism of Type IVa pilus assembly (McCallum *et al*, 2017), which involves an ATPase-driven rise and rotation of the entire nascent pilus upon subunit incorporation, is intriguingly similar to the mechanism of conjugative pilus biogenesis suggested by our model.

As mentioned in the Introduction, not all reported structures of conjugative pili feature a 1-start helical symmetry arrangement as observed in the R388 and pKpQIL structures. However, this does not exclude the possibility that such 1-start helical symmetry might be *induced* in the respective pili, as the quaternary structure of helical polymers has been demonstrated to display plasticity in response to perturbations, without significant changes to secondary and tertiary structure of the polymerising subunits (Lu *et al*, 2015; Egelman *et al*, 2015). In fact, even coexistence of geometrically distinct rotational and helical symmetries has been observed in biological filaments (Wang *et al*, 2008), including conjugative pili (Wang *et al*, 2009).

We speculate that transitions between 1-start and 5-start helical symmetry may represent one of the mechanisms by which the signal generated by contact of the pilus tip with the recipient cell surface may propagate through to the T4SS (Waksman, 2025) and to the Relaxosome in the cytoplasm to trigger DNA transfer (Williams *et al*, 2025).

The hexameric ATPase assemblies of T4SSs have been found to be directly involved in pilus subunit extraction and translocation (Kerr C Christie, 2010). In the R388 T4SS, the IMC features a hexameric assembly of TrwK_B4_ dimers (Macé *et al*, 2022), which is hypothesised to interact directly (Chetrit *et al*, 2018; Park *et al*, 2020; Khara *et al*, 2021; Yue *et al*, 2025) with the constitutively hexameric accessory ATPase TrwD_B11_ (Yeo *et al*, 2000; Hare *et al*, 2006). It is hypothesised that this interaction is required for stabilisation and coordination of the IMC protomers (Macé *et al*, 2022) and that the concerted ATPase activity of TrwK_B4_ and TrwD_B11_ is what drives pilus elaboration (Sagulenko *et al*, 2001). Consulting our model, we note that the TrwK_B4_ monomers of the inner hexameric ring are advantageously positioned to potentially make direct contact with the pilus subunits during each incorporation cycle. We hypothesise that the joint action of the tightly integrated TrwM_B3_ and ATPase subunits of the R388 IMC is what coordinates and drives unidirectional pilus elaboration. We anticipate that the methods employed in this paper will enable future efforts at achieving a detailed understanding of the mechanistic details of this crucial process underpinning bacterial conjugation and horizontal gene transfer.

## MATERIALS AND METHODS

### Strains and media

All strains used for this study are listed in Appendix Table S1.

Briefly, the *E. coli* TOP10 [StpR] strain (Thermo Fisher, Cat# C404003) was used for all expressions and was used as donor cell strain for all conjugation assays. The *E. coli* DH5α [NalR] strain (Thermo Fisher, Cat# 18265017) was used as recipient cell strain for all conjugation assays.

Strains were routinely cultured in LB-broth (Formedium, Cat# LMM0102) with shaking at 37°C. Where relevant, liquid cultures were supplemented with specific antibiotics at the following concentrations: Streptomycin (Stp) = 100 µg/mL, Nalidixic acid (Nal) = 30 µg/mL, Carbenicillin (Car) = 100 µg/mL, Kanamycin (Kan) = 50 µg/mL, Chloramphenicol (Cam) = 20 µg/mL). To achieve amber-suppressing conditions for the purpose of site-specific Bpa-incorporation, liquid cultures were supplemented with 0.2 mM 4-benzoyl-L-phenylalanine (Bpa) (Thermo Fisher, Cat# H52083.06) and expression of the *p*BpF-RS and tRNA_CUA_^opt^ encoded on the pEVOL-pBpF plasmid (Addgene, Cat# 31190) induced by addition of L-arabinose (Sigma-Aldrich, Cat# A3256) to a final concentration of 0.08%.

For colony selection and solid-support mating, LB-agar (Formedium, Cat# LMM0202) plates were used.

### Plasmids and constructs

All plasmids used and constructs generated for this study are listed in Appendix Table S2 and S3, respectively.

Plasmids used for R388 T4SS expression (pBADM11-*trwN_B1_*-*trwD_B11_*) were generated as such:

The *trwN_B1_*-*trwD_B11_* operon was cloned from wild type R388 and inserted using In-Fusion® Snap Assembly cloning (Takara, Cat# 638947) into the pBADM11 vector (Addgene, Cat# 108946) downstream of the arabinose-inducible *araBAD* promoter. The following alterations were made using In-Fusion cloning: 1) The natively occurring amber (TAG) nonsense codons of the coding sequences of *trwM_B3_*, *eex*, *trwI_B6_*, *trwE_B10_*, and *trwD_B11_* were changed to ochre (TAA) codons, 2) sequences encoding affinity tags (Flag = GSGDYKDDDDK, His = 10xH, Strep = SAWSHPQFEK) were added to the coding sequences of TrwJ_B5_, TrwI_B6_, TrwE_B10_, respectively, with all affinity tags being C-terminally placed, 3) introduction of the Thr63Cys mutation of TrwL_B2_, which has previously been shown to have no effect on conjugation and pilus biogenesis compared to wild type (Vadakkepat *et al*, 2024). Amber-substituted variants of the R388 T4SS expression plasmids were generated using In-Fusion® cloning.

The oriT-containing plasmid (pCDF-oriT-*trwA_Aux_B_D4_C_Rel_*) used for conjugation assays was generated as such:

The oriT-*trwA_Aux_B_D4_C_Rel_* locus was cloned from wild type R388 and inserted using In-Fusion® cloning into the vector pCDFDuet-1 (Addgene, Cat# 71340) downstream of the leaky T7 promoter. The existing StpR element was replaced with the KanR element from pRSFDuet-1 (Addgene, Cat# 71340) using In-Fusion® cloning.

### Expression and purification of the R388 T4SS

All expressions were carried out using the *E. coli* TOP10 strain. Vials of chemically competent *E. coli* TOP10 cells were freshly transformed with relevant plasmids, spread on LB agar plates, and kept at 37°C until individual resistant colonies emerged. Single colonies were used for inoculation of expression cultures.

For expression of the Bpa-containing variants of intact R388 T4SS, cells harbouring the pEVOL-*p*BpF plasmid and the relevant, amber-substituted R388 T4SS expression construct were cultured in 12 L LB-broth at 37°C with shaking until log phase was reached (OD_600_ = 0.5). Cultures were then supplemented with Bpa (0.2 mM) and L-arabinose (0.08%) to achieve amber-suppressing conditions and transferred to 16°C with shaking overnight.

To isolate intact R388 T4SS, cells were harvested by centrifugation (4,500 rcf, 15 min, 4°C) and the pellets (∼30 g from 12 L) resuspended in 180 mL Lysis buffer (50 mM HEPES pH 7.6, 200 mM sodium acetate, 1 mM EDTA) supplemented with protease inhibitors (Roche, Cat# 04693116001), 10 µg/mL lysozyme (Sigma-Aldrich, Cat# L6876), and 25 U/mL benzonase (Millipore, Cat# E1014). After stirring at 4°C for 15 min, the cells were lysed by multiple passages of the suspension (at 0, 10,000, 15,000, and 20,000 PSI of applied pressure) through an Emulsiflex C5 (Avestin) cooled to 4°C. The lysate was then cleared by centrifugation (27,200 rcf, 30 min, 4°C) and the membrane fraction harvested by ultracentrifugation (142,400 rcf, 45 min, 4°C). Membrane pellets were flash-frozen in liquid N_2_ and stored at -80°C until use.

To purify intact R388 T4SS, membrane pellets were manually homogenised in solubilisation buffer (50 mM HEPES pH 7.6, 200 mM sodium acetate, 0.5% *n*-dodecyl-β-D-maltopyranoside (DDM) (Anatrace, Cat# D310), 0.5% glyco-diosgenin (GDN) (Anatrace, Cat# GDN101), 0.05 mM *n*-tetradecyl-N,N-dimethylamine-N-oxide (TDAO) (Anatrace, Cat# T360), 2 mM tris(2-carboxyethyl)phosphine (TCEP) (Sigma-Aldrich, Cat# C4706), 1 mM EDTA) using a Potter-Elvehjem homogeniser (SLS, Cat# HOM3538) cooled on ice. The suspension was then incubated for 1 hr at 4°C with gentle agitation to allow for complete solubilisation. The soluble fraction was isolated by centrifugation (27,200 rcf, 20 min, 4°C) and filtered once through a 0.45 µm PES filter (Millipore, Cat# SLHPR33RS) before being loaded onto 2x 5 mL StrepTrap XT columns (Cytiva, Cat# 29401323) connected in series and equilibrated in loading buffer (50 mM HEPES pH 7.6, 200 mM sodium acetate, 0.1% GDN, 0.05 mM TDAO). A Strep affinity tag was introduced at the C-terminus of the TrwE_B10_ component (see above), which is present in 14 to 16 copies per intact complex and interacts directly with all other components of the R388 T4SS (Macé C Waksman, 2024). Our purification strategy thus ensured selective enrichment of *intact* T4SS on the column resin (Macé *et al*, 2022). After extensive washing, intact R388 T4SS was eluted using loading buffer supplemented with 50 mM biotin (Sigma-Aldrich, Cat# B4501). The eluate was concentrated by tangential flow filtration using a 100 kDa cut-off PES chip mounted on a µPulse V3 (Formulatrix) and all preparations used immediately thereafter.

### Photo-crosslinking

For *in vitro* photo-crosslinking of purified R388 T4SS, 50 uL aliquots of purified, concentrated complex were added to individual wells of Nunc MaxiSorp Flat-bottom 96-well plates (Invitrogen, Cat# 11530627) pre-washed with loading buffer and cooled on ice. These aliquots were then either left untreated or subjected to UV-irradiation. To achieve proper photo-crosslinking, the relevant 96-well plates were placed flat on ice-cold aluminium blocks carefully placed inside a CL-1000 Ultraviolet Crosslinker (UVP) fitted with 5 x 8-watt 365nm UV dual bipin discharge tubes. The plates were positioned exactly 10 cm from the tubes and the entire apparatus carefully placed on top of a Solaris SK2000 orbital shaker (Thermo Fisher, Cat# 16221591) fitted with an 18x18” platform (Thermo Fisher, Cat# 16251581) operating at 50 rpm to ensure continuous stirring.

For *in vivo* photo-crosslinking of intact R388 T4SS, donor cells producing the complex under amber-suppressing conditions were harvested and resuspended in ice-cold PBS (Gibco, Cat# 10010023) to OD_600_ = 20. The suspension was then added to Nunc 245x245x25mm BioAssay dishes (Thermo Fisher, Cat# 10489282) placed flat on ice-cold aluminium blocks carefully placed inside the CL-1000 Ultraviolet Crosslinker. The plates were then irradiated similarly to how the 96-well plates were irradiated during *in vitro* photo-crosslinking, except that the orbital shaker was operated at 25 rpm to avoid spills.

### Western blot analysis

Aliquots of purified and concentrated, intact R388 T4SS (UV-treated and untreated) were analysed by SDS-PAGE analysis using 10-well 4-12% gradient Bis-Tris gels (Invitrogen, Cat# NW04120BOX) run at room temperature (RT) for 22 min at 200 V in 1x MES running buffer (Invitrogen, Cat# B0002-02). Samples were mixed 3:1 (v/v) with 4x LDS Sample buffer (Invitrogen, Cat# B0008) freshly supplemented with 200 mM DTT (Sigma-Aldrich, Cat# D0632) and boiled at 95°C for 5 min before loading.

Gel contents were transferred onto nitrocellulose membranes (Invitrogen, Cat# IB33002) using the ‘Broad Range’ program of the iBlot3 system (Invitrogen) and membranes subjected to simultaneous blocking and blotting (see below).

For blots involving a single, HRP-conjugated antibody against either Flag-(Invitrogen, Cat# MA1-91878-HRP), His-(Invitrogen, Cat# MA1-21315-HRP), or Strep-tag (Millipore, Cat# 71591), antibody preparations were diluted 1:1,000 (v/v), 1:1,000 (v/v), or 1:4,000 (v/v), respectively, in Block-Blot buffer (1x TBS supplemented with 0.1% TWEEN® 20 (Sigma-Aldrich, Cat# P1379) and 3% (w/v) BSA (Sigma-Aldrich, Cat# A3059)). Membranes were incubated in the relevant Block-Blot buffer overnight at 4°C with shaking and then washed extensively (5x 5 min at RT with shaking in 10 mL 1xTBS supplemented with 0.1% TWEEN® 20, followed by 1x 5 min in 10 mL 1xTBS) before development using chemiluminescent substrate (Thermo Fisher, Cat# 34577). The blots were visualised and photographed using a ChemiDoc MP Imaging System (Bio-Rad).

For blots involving both primary unconjugated and secondary HRP-conjugated antibodies, membranes were first incubated in the relevant *primary* Block-Blot buffer (containing 1:2,000 (v/v) Rabbit anti-OmpA IgG (Thermo Fisher, Cat# PA5-144390)) overnight at 4°C with shaking and then washed extensively (see above) before being incubated in the relevant *secondary* Block-Blot buffer (containing 1:50,000 (v/v) Goat anti-Rabbit HRP-conjugated IgG (abcam, Cat# ab205718)) for 1 hr at RT with shaking.

Washing, development, and visualisation was then carried out as described above.

### Mass spectrometry sample preparation

Duplicate Coomassie-stained gels were prepared in parallel with all gels used for Western blot analysis. These duplicate gels were used for mass spectrometry analysis. For preparation of in-gel digested samples for MS analysis, individual bands from Coomassie stained gels were carefully dissected and destained using 30% ethanol until all visible stain was removed. Gel bands were then diced and dehydrated using 100% ACN. Reduction and alkylation of band contents was achieved by rehydrating the gel band in 1M Tris pH 8.5 supplemented with 5 mM TCEP and 20 mM CAA and incubating for 5 mins at 70°C with 650 rpm shaking. The gel band was then extensively washed (3x 1 min at 70°C with 650 rpm shaking) in 50% ethanol in 50 mM Tris pH 8.5 and once again dehydrated using 100% ACN.

For single-digest using Chymotrypsin, each gel band was incubated in 100 µL of 100 mM Tris pH 8.0, 10 mM CaCl_2_ with 2.5 µL 1 µg/µL Chymotrypsin (freshly dissolved in 1 mM HCl) and incubated for 16 hrs at 37°C with 650 rpm shaking.

For double-digest using Trypsin and Chymotrypsin in sequence, each gel band was first incubated in 100 µL of 100 mM Tris pH 8.5 with 10 µL 100 ng/µL Trypsin (dissolved in 50 mM acetic acid) and incubated for 16 hrs at 37°C with 650 rpm shaking. Samples were then supplemented with 10 mM CaCl_2_ and 2.5 µL 1 µg/µL Chymotrypsin (freshly dissolved in 1 mM HCl) and incubated an additional 4 hrs at 37°C with 650 rpm shaking. Enzymatic digestion was quenched by addition of 10 uL 10% TFA and the quenched digest reaction transferred to a clean tube. The gel piece was incubated in 50 µL 100% ACN at RT for 1 min and this wash mixed with the quenched digest. The tubes containing the digest reactions were transferred to a SpeedVac DNA 130 (Thermo Fisher, Cat# DNA130-230) and run for 2 hrs at 37°C until all solvent was evaporated. Immediately prior to MS analysis, the dehydrated samples were rehydrated and acidified in 50 uL 0.1% TFA.

### Mass spectrometry analysis

Acidic samples of rehydrated peptides were separated on a µPAC Neo Plus HPLC Column (50 cm, pore size 100-300 Å) using a micro-LC UltiMate 3000 high-performance liquid chromatography system. Peptides were eluted over a linear gradient from 1 to 35% solvent B (80% ACN) for 24 min at a flow rate of 50 µL/min. MS/MS data were collected in data-dependent mode using an in-line Orbitrap Eclipse instrument (Thermo Fisher) equipped with an H-ESI source. MS1 scans were acquired in the Orbitrap analyser at a resolution of 120,000 using a scan range of either *m/z* 375-1,500 or *m/z* 350-2,000. Precursor ions were isolated using the quadropole (isolation window 1.4 m/z), including only charge states 2-7. After HCD with a normalised collision energy of 30%, MS2 scans were acquired in the Orbitrap analyser at a resolution of 30,000 using a minimal intensity threshold of 50,000 and dynamic exclusion time set to 12 s.

For identification of peptides, the resulting MS data were analysed and validated using MSFragger (Kong *et al*, 2017) and Philosopher (da Veiga Leprevost *et al*, 2020) as implemented in the FragPipe (v23.0) analysis suite (available at www.fragpipe.nesvilab.org). Briefly, a search database was generated by combining the *E. coli* K12 proteome (Proteome ID: UP000000625) with fasta files containing the amino acid sequences of all the R388 elements encoded on the employed plasmids. Positions corresponding to amber-substituted codons were changed to ‘X’, and ‘X’ introduced as a fixed modification (Mass Delta = 251.09464) to accurately reflect the mass of an incorporated Bpa residue.

MS1 and MS2 spectra were visualised and manually evaluated using FreeStyle (Thermo Fisher). Isotope simulations were generated and corresponding ion chromatograms extracted using the built-in Isotope Simulation tool. Individual peaks were identified and analysed using the built-in Genesis Peak Detection Algorithm using default parameters. MS1 spectra were visualised using FreeStyle, while MS2 spectra and corresponding ion tables were generated and visualised using the PDV Viewer as implemented in FragPipe.

### Conjugation assays

Vials of electrocompetent *E. coli* TOP10 cells [StpR] were freshly transformed with *p*BpF-pEVOL [CamR], pCDF-oriT-*trwA_Aux_B_D4_C_Rel_* [KanR], and an amber-less *or* relevant amber-substituted variant of the pBADM11-*trwN_B1_*-*trwD_B11_*plasmid [CarR] encoding the R388 T4SS. Transformations were spread on LB agar plates containing 100/50/20 µg/mL Car/Kan/Cam and kept at 37°C until individual, resistant colonies emerged. Single colonies were used for inoculation of selective LB-broth and cultured overnight at 37°C with shaking to generate donor cell stock cultures.

Recipient cells (*E. coli* DH5α strain) [NalR] were freshly streaked from frozen glycerol stock onto LB agar plates containing 30 µg/mL Nal and kept at 37°C until individual, resistant colonies emerged. Single colonies were used for inoculation of selective LB-broth and cultured overnight at 37°C with shaking to generate recipient cell stock cultures.

Stock cultures of donor and recipient cells were used for inoculation of separate flasks of 200 mL selective LB-broth to a starting OD_600_ of 0.05. Cultures were kept at 37°C with shaking until all donor cell cultures reached an OD_600_ of 0.5-0.6. The donor cell cultures were then supplemented with 0.2 mM Bpa (80 uL of 0.5 M stock in 1M NaOH) and 0.08% L-arabinose (800 uL of 20%, sterile-filtered stock) and all cultures transferred to 16°C with shaking overnight to allow for amber-suppressed expression of intact R388 T4SS.

Donor cell masses corresponding to 60 mL OD_600_ 1.0 culture (= 60 OD-units (ODUs)) were harvested and resuspended in 30 mL ice-cold 1x PBS (to OD_600_ 2.0). 15 mL of each donor cell suspension (30 ODUs) was either kept on ice (untreated) or transferred to individual Petri dishes kept on ice-cold aluminium blocks carefully placed inside a CL-1000 Ultraviolet Crosslinker (UVP) for UV-treatment. The plates were positioned exactly 10 cm from the tubes and the entire apparatus carefully placed on top of an orbital shaker (see above) operating at 50 rpm to ensure continuous stirring of the donor cell suspension. The plates were then subjected to 365 nm wavelength near-UV irradiation for 30 minutes (corresponding to a dose of 3.6 J/cm^2^) with constant cooling and stirring. 1.0 ODU of untreated and UV-treated donor cell populations were then pelleted by centrifugation (5,000 rcf, 2 min, 4°C), resuspended in 0.5 mL fresh LB-broth, and kept unstirred at 37°C for 2 hrs to allow for recovery. Remaining donor cells were pelleted, flash-frozen, and kept at -80°C for SDS-PAGE and Western blot analysis of whole cell extracts. Whole cell extracts were prepared by resuspending individual pellets (∼50 mg, corresponding to 10 ODUs) in 200 uL 1x Bugbuster® protein Extraction Reagent (Millipore, Cat# 70584-3) supplemented with 0.2 mg/mL lysozyme and 50 U/mL benzonase and incubating the suspension at RT with shaking until the suspension became clear. Amounts of clear lysate corresponding to 1.0 ODU were analysed in each lane during SDS-PAGE.

For solid-support mating, 1.0 ODU of recovered donor cells and 1.0 ODU of freshly cultivated recipient cells were mixed and pelleted by centrifugation (5,000 rcf, 2 min, RT). The resulting pellet was resuspended in 100 uL LB-broth and pipetted onto sterile nitrocellulose filter papers (Millipore, Cat# HAWP02500) placed directly on semi-dry LB agar plates. The plates were placed upside down at 37°C for 2 hrs to allow for mating to occur. The mating pellets were harvested by carefully removing the filter papers and washing them twice with 0.5 mL LB-broth. Serial dilutions of the resulting mating suspension were then plated on either donor (D) cell selective LB agar plates (100/50 µg/mL Stp/Kan) or transconjugant (T) cell selective LB agar plates (30/50 µg/mL Nal/Kan) and kept at 37°C for at least 16 hrs until resistant colonies emerged.

To calculate the number of colony forming units (CFUs) for each plate, the following formula was used: #CFUs = [(#colonies counted * 10^#dilutions performed^)/100 uL]*1,000 uL, which simplifies to #CFUs = (x*10^y^)*10, with x = #colonies counted and y = #dilutions performed. The ratio of T cells and D cells (T/D) was used as a measure of conjugation efficiency.

### Quantification and statistical analysis

For the conjugation assays, the data points represent measurements of conjugation efficiencies (calculated as described above) determined from independent experiments. At least three independent experiments (n ≥ 3) were performed for each condition.

A Bonferroni’s multiple comparisons test comparing preselected pairs of means (as implemented in GraphPad Prism 10) was used to determine statistical significance (indicated in figure panels: Asterisks denoting statistical significance; ’ns’ denoting ’no significance’) by computing multiplicity-adjusted P-values (reported explicitly in figure legends and main text).

### Structural modelling and structure prediction

ChimeraX (Meng *et al*, 2023) was used for visualisation and modelling. We used the following published structures: Hexameric composite model of the R388 IMC (IMC model) (PDB: 7O41; EMD-12715) (Macé *et al*, 2022); R388 conjugative pilus model (pilus model) (PDB: 8S6H; EMD-19758) (Vadakkepat *et al*, 2024), Extended inner membrane complex (IMC) protomer structure (R388 IMC protomer) (PDB: 8RTB; EMD-19485), Stalk-Arches-IMC structure (R388 Stalk+IMC) (PDB: 8RTD; EMD-19488) (Macé C Waksman, 2024). Additional structures used for figure preparation: OMCC O-layer/I-layer (PDB: 7O3J/7O3T) (Macé *et al*, 2022). AlphaFold3 (AF3) (Abramson *et al*, 2024) (available at www.alphafoldserver.com) was used for structure prediction of the binary TrwL_B2_-TrwM_B3_ and TraM_B2_-TraA_B3_ complexes. The complete amino acid sequences of the VirB3 moieties (residues 1-104 for TrwM_B3_, residues 1-105 for TraA_B3_) were used, while only the amino acid sequences corresponding to the mature version of the VirB2 components (residues 44-112 for TrwL_B2_, residues 28-97 for TraM_B2_), in which the signal peptides have been removed, was used for modelling.

Our composite model of the pilus biogenesis state(s) of the R388 T4SS was generated by manually docking the convex end of the pilus model onto the concave IMC model. The pilus model was positioned relative to the IMC model by defining a mass-weighted, longitudinal axis of the pilus structure and aligning this to the normal of a mass-weighted transverse plane of the IMC (generated from the coordinates of the six modelled copies of TrwM_B3_). This *horizontally* aligned pilus model was appropriately positioned *vertically* based on general shape complementarity and avoiding steric clashes between incorporated TrwL_B2_ subunits and the subunits of the relevant IMC protomers.

The ∼2.6 Å rise and ∼6° rotation of the entire pilus per incorporation cycle was simulated using ChimeraX by applying the following geometric operation: *turn [plane normal] -c.1 model [pilus] center [plane normal]; move [plane normal] 2.c2 model [pilus model]*, which results in an exactly 6.1° left-handed (*CW* as seen from above) rotation around the aligned axes, and a 2.62 Å rise (towards the viewer as seen from above) along these axes.

At the end of select incorporation cycles, pairs of the last to be incorporated TrwL_B2_ subunit model and the corresponding TrwM_B3_ subunit model were saved as separate composite models and superimposed on each other along their TrwM_B3_ moieties (see Fig EV7). Cɑ RMSD calculations were then carried out for the respective TrwL_B2_ moieties using ChimeraX to confirm geometric consistency in the TrwL_B2_-TrwM_B3_ association throughout arbitrarily selected incorporation cycles.

The ‘Binding state’ configuration of the TrwL_B2_-TrwM_B3_ interaction during pilus subunit incorporation was predicted by rigid-body alignment of the TrwL_B2_-TrwM_B3_ AF3-prediction onto the copies of TrwM_B3_ in the IMC model using the ChimeraX ‘matchmaker’ tool.

The ‘Recruitment state’ configuration of the TrwL_B2_-TrwM_B3_ interaction during pilus subunit incorporation was based on the (experimentally determined) equivalent state of the TraM_B2_-TraA_B3_ interaction of the pKM101 IMC protomer (see below).

Briefly, the TrwL_B2_ moiety of the AF3-predicted TrwL_B2_-TrwM_B3_ complex was manually truncated to match the equivalent sequence stretch represented by distinct density in the relevant *pKM101* IMC protomer map. This truncated TrwL_B2_ moiety was then fitted into the equivalent density of the published map of the *R388* IMC protomer.

### Expression and purification of the pKM101 T4SS

In order to purify the pKM101 T4SS, the plasmid pBADM11_*traL_B1_*-*traF_B10_CStrep*_*rbstraG_B11_*-*traJ_D4_*(a clone shown to mediate conjugation) was modified to delete the *traJ_D4_* gene while simultaneously tagging TraG_B11_ with a C-terminal His tag using an inverse-PCR strategy followed by In-Fusion cloning. This construct, named as the pBADM11-pKM101T4SS-ΔD4 construct was used for all subsequent purification of the pKM101 T4SS described below.

To purify the pKM101 T4SS complex, the plasmid pBADM11-pKM101T4SS-ΔD4 was introduced into *E. coli* TOP10 cells via electroporation. Transformants were selected by plating on Luria-Bertani (LB) agar supplemented with 100 μg/mL carbenicillin. To minimise leaky expression from the arabinose-inducible *pBAD* promoter, which can adversely affect cell growth, 0.5% (w/v) glucose was included in both the LB agar plates and the initial liquid LB cultures. An overnight culture (1%) of the freshly transformed *E. coli* TOP10 cells harbouring the pBADM11-pKM101T4SS-ΔD4 plasmid was inoculated into 12 litres of LB medium containing 100 μg/mL carbenicillin. Cultures were incubated at 37°C with agitation at 180 rpm until reaching an OD₆₀₀ of 0.5, at which point protein expression was induced with 0.08% (w/v) L-arabinose. Induction proceeded overnight under the same growth conditions. Cells were harvested by centrifugation at 5,000 × *g* for 30 minutes and resuspended in lysis buffer comprising 50 mM HEPES (pH 7.6), 200 mM sodium acetate, and 1 mM EDTA (pH 8.0), supplemented with four protease inhibitor cocktail tablets, 1 mg/mL lysozyme, and 1 μg/mL Benzonase nuclease. Cell lysis was achieved using an EmulsiFlex-C3 high-pressure homogeniser. Cellular debris and inclusion bodies were removed by centrifugation at 26,712 × *g* for 30 minutes. Membranes were subsequently isolated by ultracentrifugation at 95,834 × *g* for 45 minutes. The resulting membrane pellet was homogenised using a Potter-Elvehjem homogeniser in solubilisation buffer consisting of 50 mM HEPES (pH 7.6), 200 mM sodium acetate, 1 mM EDTA (pH 8.0), 0.5% (w/v) DDM, 0.5% GDN, 0.05 mM TDAO, and 2 mM TCEP. The homogenised membrane suspension was incubated at 4°C for 1 hour on a rotary shaker to enable detergent-mediated solubilisation of the pKM101 T4SS complex. Insoluble material was removed by a second ultracentrifugation at 95,834 × *g* for 20 minutes. The clarified supernatant was loaded onto a 5 mL StrepTrap HP column (Cytiva) pre-equilibrated with buffer A (50 mM HEPES, pH 7.6; 200 mM sodium acetate; 0.1% GDN; 0.05 mM TDAO) at a flow rate of 1 mL/min. After extensive washing with buffer A, the protein complex was eluted with buffer A supplemented with 2 mM desthiobiotin.

Following Strep elution of the pKM101 T4SS complex, samples were processed on sucrose gradients prepared in buffer D (50 mM HEPES, pH 7.6; 200 mM sodium acetate; 0.1% GDN; 0.05 mM TDAO). Gradient fractions were collected and assessed for protein composition and purity by SDS-PAGE. The presence of intact pKM101 T4SS assemblies was verified via negative-stain electron microscopy (NSEM) using 2% uranyl acetate as the stain. Fractions containing intact pKM101 T4SS complexes were subsequently buffer-exchanged to remove sucrose using overnight dialysis in 50 mM HEPES (pH 7.6) and 200 mM sodium acetate (with detergents), employing 6–8 kDa molecular weight cut-off D-Tube Dialysers Maxi (Merck, Cat# 71509-M). Following dialysis, samples were centrifuged at 15,000 × *g* for 15 minutes to remove protein aggregates prior to cryo-electron microscopy (cryo-EM) grid preparation.

### Cryo-EM grid preparation and data acquisition

3 μL of a sample showing intact pKM101 T4SS in NSEM and at a concentration of 0.03 mg/ml was used for cryo-EM grid preparation. The sample was applied to UltrAuFoil Au200 R 2/2 grids (Quantifoil, Germany) coated with graphene oxide (Naydenova *et al*, 2019). Prior to application, the grids were negatively glow-discharged at 25 mA for 60 seconds using a PELCO easiGlow device (Ted Pella). Following sample application, the grids were incubated for 20–30 seconds in a Vitrobot Mark IV (Thermo Fisher Scientific) chamber maintained at 4 °C and 100% relative humidity, after which they were blotted for 16 s and vitrified in liquid ethane.

The cryo-EM data were collected at the ISMB Birkbeck EM facility using a Titan Krios transmission electron microscope (Thermo Fisher Scientific, USA) operated at an accelerating voltage of 300 keV. The microscope was equipped with a BioQuantum energy filter (Gatan) with a 20 eV slit width and a K3 direct electron detector (Gatan) operating in super-resolution mode. Data were collected at a nominal magnification of 105,000X, corresponding to a calibrated pixel size of 0.828 Å. The electron dose rate was set to 13.7 electrons per pixel per second, with a total exposure of 50.3 electrons per Å² fractionated across 50 frames. Automated data acquisition was carried out using the EPU software, with a defocus range of −0.9 μm to −2.4 μm. A total of 6 individual data collection sessions totalling 170,880 movies were collected.

### Image processing of the pKM101 T4SS

All the datasets were first individually processed in Relion 4.0 (Scheres, 2012) to carry out motion correction using MOTIONCOR2 (Zheng *et al*, 2017). The resulting micrographs were imported into CRYOSPARC v4 (Punjani *et al*, 2017) for subsequent processing. Following CTF estimation using the Patch CTF implementation in CRYOSPARC, micrographs with devitrified ice, poor particle distribution and poor CTF-fit were manually removed to yield good micrographs on which particle picking was performed using the CRYOSPARC Blob picker using an elliptical blob. By optimizing the picking diameter, particles focussed and centred on the IMC could be picked up successfully using this process. All the picked particles were inspected using the inspect particle pick program, extracted using a box size of 400 Å and subjected to multiple rounds of 2D classification to clean the particle dataset.

### Image processing of the pKM101 T4SS Stalk+IMC

An initial twofold binned particle subset consisting of 251,803 particles was processed. Homogeneous refinement using the low-pass filtered (50 Å) R388 Stalk+IMC map as an initial reference yielded a reconstruction at 8.43 Å resolution. Iterative homogeneous refinement with increasingly stringent low-pass filtering improved this to 8.05 Å, which was subsequently refined using multiple rounds of non-uniform refinement to yield a map at 4.73 Å resolution. Homogeneous refinement of the unbinned particle dataset, employing adjusted masking parameters (inner radius = 0.85; outer radius = 0.95), and further non-uniform refinement, resulted in a final reconstruction of the Stalk+IMC complex at 3.96 Å. Local refinement using a Stalk+IMC mask improved the resolution to 3.86 Å. Focused local refinement using an IMC protomer mask yielded a reconstruction at 3.56 Å resolution.

### Model building of the pKM101 T4SS IMC protomer

Reported map resolutions were calculated using local resolution estimation (LocRes), which are marginally higher than the global resolution estimates obtained from CRYOSPARC refinement jobs. Model building was guided by the 3.60 Å local refinement map of the IMC protomer.

The unsharpened density map was subjected to post-processing to enhance interpretability. Two sharpening strategies were employed: (a) the tight-target mode in DeepEMhancer (Sanchez-Garcia *et al*, 2021), which facilitated improved map clarity and was particularly advantageous for downstream integration with automated model-building tools such as ModelAngelo (Jamali *et al*, 2024); and (b) PHENIX Autosharpen (Terwilliger *et al*, 2018), which was used in parallel to enhance structural features for manual model building. Models generated via automated pipelines were subsequently adjusted manually in Coot (Emsley C Cowtan, 2004) and refined against the PHENIX autosharpened maps using PHENIX.

Due to the dynamic nature of the pKM101 T4SS IMC, well-resolved density was observed for only three protomers. Of these, only two displayed continuous density, with protomer 1 yielding superior map quality. Model building efforts therefore focused on this protomer, comprising two copies of TraB_B4_, one TraM_B3_, and four N-terminal helices of TraE_B8_ (residues 5–62). Model building of the IMC protomer was also challenging owing to the severe preferential orientation exhibited by the outer extremities of the IMC protomer, especially the TraE_B8_ helices, making the interpretation of these densities and model building of these regions extremely challenging. Hence, instead of using fully automated model building tools like ModelAngelo, we chose to proceed with using semi-automated model building methods. This involved generating a homology model of the IMC protomer by carrying out a one-on-one threading of the pKM101 sequences (2×TraB_B4_, 1×TraA_B3_, 4×TraE_B8_) onto the R388 IMC protomer structure. This model was docked into the DeepEMhancer-sharpened IMC map in Chimera, followed by RosettaCM-based (Song *et al*, 2013) semi-automated modelling and iterative manual adjustments in Coot against the PHENIX autosharpened map.

In order to build the TraM_B2_ helix-loop-helix region (amino acids 45-89) in the IMC protomer, we attempted building this region in all four possible combinations ((i) TraM_B2_ loop facing the cytoplasm, (ii) its flipped version, (iii) TraM_B2_ loop facing the periplasm, and (iv) its flipped version)). Fortunately, the presence of amino acid residues with bulky side chains at the turn allowed us to place the TraM_B2_ subunit with high confidence. After the initial placement of the TraM_B2_ chain was worked out, the chain corresponding to TraM_B2_ was manually adjusted in Coot against the PHENIX autosharpened map to optimise for main chain and side chain fitting. The manually adjusted model of the IMC protomer comprising of 2×TraB_B4_, 1×TraM_B3_, and 4×TraW_B8_ was refined against the PHENIX autosharpened map using real space refinement in PHENIX followed by optimization of Molprobity (Chen *et al*, 2009) parameters to finalise the IMC protomer model. The Coot-adjusted chain corresponding to TraM_B2_ (for which PHENIX real space refinement resulted in a much less plausible main and side chain density fitting) was then added separately to the finalised IMC protomer model.

## DATA AVAILABILITY

• The electrostatic potential maps for the pKM101 ’IMC+Stalk’ and the pKM101 ’IMC protomer’ were deposited in the EMDB under accession codes: EMD-58169 and EMD-58133, respectively.

• Atomic coordinates for the pKM101 ’IMC protomer’ were deposited in the Protein DataBank under accession code: 30XS.

## AUTHOR CONTRIBUTIONS

G.W., M.B., A.V, and K.T. conceptualized the research project and planned the experiments.

M.B. carried out all R388 T4SS expressions, purifications, and sample preparations for photo-crosslinking, Western blot, and conjugation assays, performed the experiments, and analysed the data.

A.V. carried out all pKM101 T4SS expressions, purifications, and sample preparations for cryo-EM analysis, and analysed the data.

M.B. performed all R388 T4SS structural modelling and all R388 and pKM101 T4SS structure predictions.

A.V. built, refined, and validated the pKM101 T4SS structural models.

M.B. and A.V. carried out all molecular cloning.

M.B. prepared all samples for mass spectrometry.

R.Z. and M.B. collected the mass spectrometry data.

M.B. wrote the first draft of the paper and prepared all the figures together with G.W.

G.W. and K.T. secured the funding for the project.

All authors contributed to the revision of the manuscript.

## ACKNOWLEDGEMENTS

This work was supported by MRC grant 01827 (awarded to G.W. and K.T.) and Wellcome grant 217089 (awarded to G.W.). We thank the ISMB EM facility at Birkbeck College, University of London and the University College London Mass Spectrometry Science Technology Platform. We thank D. Houldershaw for IT support.

