## Supplementary tables and figures for "*In vivo* crosslinking and cryo-EM analyses suggest a revised model of conjugative pilus biogenesis"

Appendix Table S1

Strains used in this study:

| **Strain** | **Description** | **Reference** |
|---|---|---|
| *E. coli* TOP10 | F- mcrA Δ(mrr-hsdRMS-mcrBC) Φ80lacZΔM15 Δ lacX74 recA1 araD139 Δ(araleu)7697 galU galK rpsL (StrR) endA1 nupG.<br>Used for cloning, expressions, and conjugation assays. | Thermo Fisher, Cat# C404003 |
| *E. coli* DH5α | $F^-$ φ80*lac*ZΔM15 Δ(*lac*ZYA-*arg*F)U169 *rec*A1 *end*A1 *hsd*R17($r_K^-$, $m_K^+$) *pho*A *sup*E44 $\lambda^-$*thi*-1 *gyr*A96 *rel*A1<br>Used for conjugation assays. | Thermo Fisher, Cat# 18265017 |

Appendix Table S2

Plasmids used in this study:

| **Plasmid** | **Description and Reference** | **Source** |
|---|---|---|
| R388 | Wild-type T4SS conjugative plasmid of *E. coli* belonging to IncW incompatibility group, Trimethoprim resistance | Lab material (pR388-47) |
| pKM101 | Wild-type T4SS conjugative plasmid of *E. coli* belonging to IncN incompatibility group, historically significant for its use in mutagenicity testing, particularly in the Ames test, where it enhances the sensitivity of certain *Salmonella* tester strains to mutagens by increasing error-prone DNA repair processes, Ampicillin resistance | Lab material (pKM101-3) |
| pBADM11 | https://www.addgene.org/108946/ | Addgene, Cat# 108946 |
| pCDFDuet-1 | https://www.addgene.org/vector-database/2080/ | Addgene, Cat# 71340 |
| pRSFDuet-1 | https://www.addgene.org/vector-database/4005/ | Addgene, Cat# 71340 |
| pEVOL-pBpF | https://www.addgene.org/31190/ | Addgene, Cat# 31190 |
| pBADM11_*traL*$_{B1}$-*traF*$_{B10}$CStrep_rbs*traG*$_{B11}$-*traJ*$_{D4}$ | Plasmid encoding the pKM101 T4SS including ATPase TraG$_{B11}$ and coupling protein TraJ$_{D4}$ with a C-terminal Strep tag on TraF$_{B10}$. | Lab material (pKM101-68) |

Appendix Table S3

Constructs generated for this study:

| **Construct** | **Description and Reference** | **Source** |
|---|---|---|
| pBADM11-*trwN*$_{B1}$-*trwD*$_{B11}$ | Plasmid encoding the R388 T4SS where the following alterations have been made:<br>1) The natively occurring amber (TAG) nonsense codons of the coding sequences of *trwM*$_{B3}$, *eex*, *trwI*$_{B6}$, *trwE*$_{B10}$, and *trwD*$_{B11}$ have been changed to ochre (TAA) codons,<br>2) sequences encoding affinity tags (Flag = GSGDYKDDDDK, His = 10xH, Strep = SAWSHPQFEK) have been added to the coding sequences of TrwJ$_{B5}$, TrwI$_{B6}$, TrwE$_{B10}$, respectively, with all affinity tags being C-terminally placed,<br>3) the T63C mutation of *trwL*$_{B2}$, which has previously been shown to have no effect on conjugation and pilus biogenesis compared to wild type (Vadakkepat *et al*, 2024), has been introduced. Amber-substituted variants of this R388 T4SS expression plasmid were generated using In-Fusion cloning. | This study |
| pCDF-oriT-*trwA*$_{Aux}$*B*$_{D4}$*C*$_{Rel}$ | oriT-containing plasmid used for conjugation assays. | This study |
| pBADM11-pKM101T4SS-ΔD4: *traF*$_{B10}$CStrep_rbs*traG*$_{B11}$-CHis-Δ*traJ*$_{D4}$ | Plasmid encoding the pKM101 T4SS where TraJ$_{D4}$ was deleted using inverse PCR while simultaneously incorporating a His-tag at the C-terminal of TraG$_{B11}$. A C-terminal Strep-tag was introduced on TraF$_{B10}$. | This study |

Appendix Table S4

Key figures of cryo-EM data collection, processing, refinement, and validation.

| Structures generated in this study | IMC Protomer | IMC+Stalk complex |
|---|---|---|
| PDB entry | 30XS | --- |
| Refinement | | |
| EMDB corresponding | EMD-58133 | EMD-58169 |
| Initial model used | *De novo* | |
| Model resolution (Å) | 3.5 | |
| FSC threshold | 0.5 | |
| CC Model vs. Data (mask) | 0.59 | |
| Model composition | | |
| Nonhydrogen atoms | 15144 | |
| Protein residues | 1882 | |
| Ligands | --- | |
| B factors (Å$^2$) | | |
| Protein | 2 (30.0 – 1.02) | |
| Ligand | --- | |
| R.m.s. deviations | | |
| Bond lengths (Å) | 0.003 | |
| Bond angles (°) | 0.705 | |
| Validation | | |
| MolProbity score | 2.61 | |
| Clashscore | 14.79 | |
| Rotamer outliers (%) | 3.45 | |
| Ramachandran plot | | |
| Favored (%) | 90.96 | |
| Allowed (%) | 8.06 | |
| Disallowed (%) | 0.97 | |

FIGURE EV1

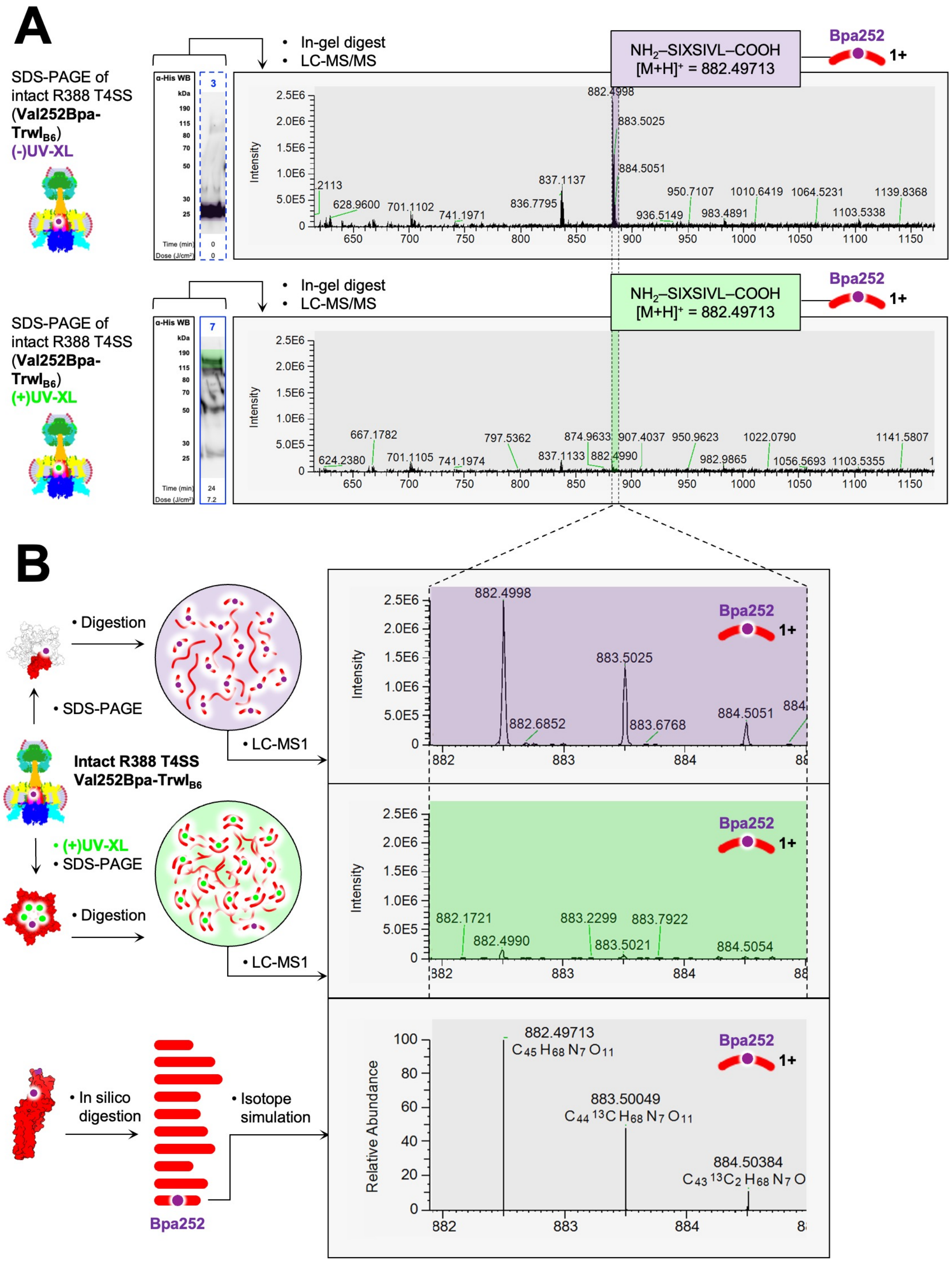
A
SDS-PAGE of intact R388 T4SS (Val252Bpa-TrwI$_{B6}$) (-)UV-XL
In-gel digest
LC-MS/MS
NH$_2$–SIXSIVL–COOH [M+H]$^+$ = 882.49713
Bpa252
1+
Intensity
SDS-PAGE of intact R388 T4SS (Val252Bpa-TrwI$_{B6}$) (+)UV-XL
B
Digestion
SDS-PAGE
LC-MS1
Intact R388 T4SS Val252Bpa-TrwI$_{B6}$
(+)UV-XL
In silico digestion
Isotope simulation
Relative Abundance
882.49713
C45 H68 N7 O11
883.50049
C44 13C H68 N7 O11
884.50384
Bpa252

**Figure EV1**: *LC-MS/MS analysis confirming site-specific Bpa-incorporation and photo-crosslinking of the R388 T4SS Val252Bpa-TrwI$_{B6}$ variant.*

**A)** Representative precursor (MS1) spectra obtained from LC-MS/MS analyses of in-gel digestions of dissected gel bands from SDS-PAGE analysis of specific samples of untreated (purple) and UV-treated (green) complex shown in Fig 2C. Based on *in silico* chymotrypsin digestion of TrwI$_{B6}$, we determined the exact mass of the doubly ($[M+2H]^{2+}$) charged species of the proteolytic peptide $NH_2$-SIXSIVL-COOH (spanning sequence positions 250-256 and featuring the expected Val (V) to Bpa (X) substitution at position 252). Automated peak detection identified peaks corresponding to these masses with retention times of the corresponding ion chromatograms being similar (1,641.42 s and 1,641.33 s, respectively) for the untreated and UV-treated samples (see Fig EV2).

**B)** Magnified views of selected peaks of the mass spectra shown in panel A). The observed isotopic envelopes fit with the simulated isotopic envelope of the singly ($[M+H]^{+}$) charged species of the $NH_2$-SIXSIVL-COOH peptide. Selective, relative depletion (quantified in Fig EV2) of this Bpa-containing *mono*peptide in the UV-treated sample (green) relative to the untreated sample (purple) is evident.

FIGURE EV2

A B C D

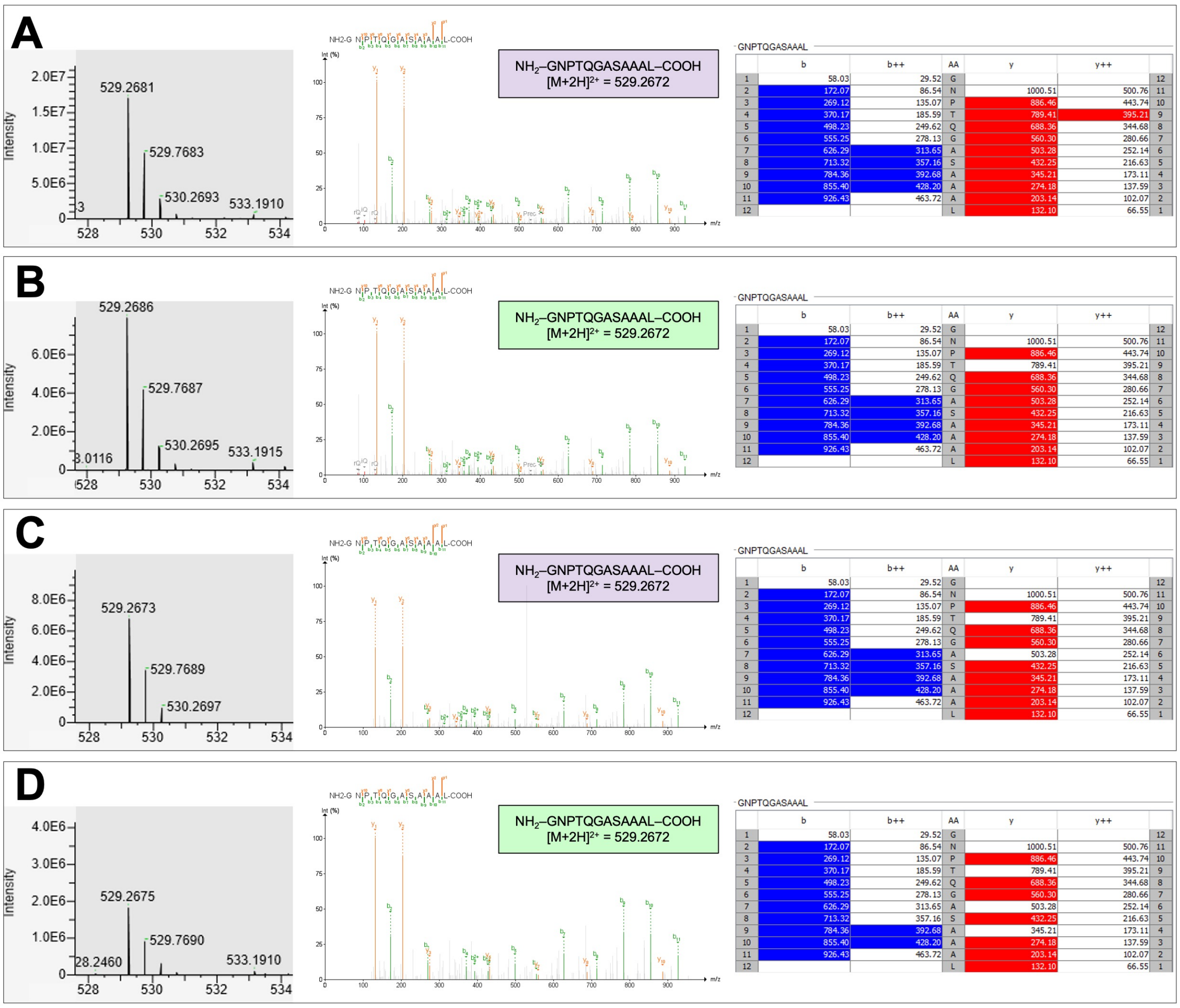

E

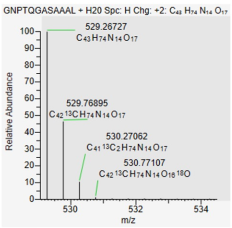

F

| Sample variant | Identity | Base Peak | Peak Area | Peak Height | RT (sec) |
|---|---|---|---|---|---|
| V252X | Control (-)UV | 529.27 | 2,106,126.44 | 17,177,340.00 | 768.68 |
| V252X | Control (+)UV | 529.27 | 2,528,231.40 | 8,228,330.00 | 775.98 |
| F136X | Control (-)UV | 529.27 | 530,612.94 | 7,067,163.00 | 642.66 |
| F136X | Control (+)UV | 529.27 | 139,688.58 | 1,822,730.62 | 643.07 |
| V252X | Bpa-pep. (-)UV | 882.50 | 982,396.08 | 2,544,745.25 | 1,641.42 |
| V252X | Bpa-pep. (+)UV | 882.50 | 162,734.52 | 205,015.16 | 1,641.33 |
| F136X | Bpa-pep. (-)UV | 1,308.54 | 207,350.33 | 1,053,232.50 | 1,430.64 |
| F136X | Bpa-pep. (+)UV | 1,308.55 | 18,318.68 | 32,267.86 | 1,427.85 |

G

| Description | Sample variant | Analysis | | Results |
|---|---|---|---|---|
| Ratio Control (∝ Total TrwI/VirB6) | V252X | Control (-)UV : (+)UV | = | 0.83 |
| Ratio Control (∝ Total TrwI/VirB6) | F136X | Control (-)UV : (+)UV | = | 3.80 |
| Ratio Bpa-pep. | V252X | Bpa-pep. (-)UV : (+)UV | = | 6.04 |
| Ratio Bpa-pep. | F136X | Bpa-pep. (-)UV : (+)UV | = | 11.32 |
| Norm. Ratio Bpa-pep. | V252X | (Bpa-pep. (-)UV : (+)UV) : (Control (-)UV : (+)UV) | = | 7.25 |
| Norm. Ratio Bpa-pep. | F136X | (Bpa-pep. (-)UV : (+)UV) : (Control (-)UV : (+)UV) | = | 2.98 |
| Relative % Decrease (+)UV vs. (-)UV | V252X | ((1/[(Bpa-pep. (-)UV : (+)UV) : (Control (-)UV : (+)UV)])-1)*100 | = | -86.2 |
| Relative % Decrease (+)UV vs. (-)UV | F136X | ((1/[(Bpa-pep. (-)UV : (+)UV) : (Control (-)UV : (+)UV)])-1)*100 | = | -66.4 |

**Figure EV2**: *Demonstration of normalised, relative, selective depletion of Bpa-containing monopeptide in proteolytic digests of UV-treated samples.*
**A-D)** Representative precursor (MS1) spectra (left), fragmentation (MS2) spectra (middle), and ion tables (right) from LC-MS/MS analyses of in-gel digestions of A) the untreated Val252Bpa-$TrwI_{B6}$ variant, B) the UV-treated Val252Bpa-$TrwI_{B6}$ variant, C) the untreated Phe136Bpa-$TrwI_{B6}$ variant, and D) the UV-treated Phe136Bpa-$TrwI_{B6}$ variant. Peaks corresponding to the doubly ($[M+2H]^{2+}$) charged species of the displayed proteolytic peptide ($NH_2$-GNPTQGASAAAL-COOH) were automatically picked and used as proxies for quantification of total amount of $TrwI_{B6}$ in the analysed samples.
**E)** Simulated isotopic envelope of the doubly ($[M+2H]^{2+}$) charged $NH_2$-GNPTQGASAAAL-COOH peptide.
**F)** Table containing relevant parameters (Base Peak (*m/z)*, Peak Area, Peak Height, Retention Time (RT) of the corresponding ion chromatograms) for the 'Control' peaks (representative spectra shown in panels A-D) and the peaks ('Bpa-pep.') corresponding to Bpa-containing peptides (representative spectra shown in Fig EV1 and 4A) that were picked by automated peak detection (see Materials and Methods) in the LC-MS/MS data acquired from the indicated samples.
**G)** Calculations of the 'Relative % Decrease' of Bpa-containing *mono*peptide (normalised to the corresponding 'Control' Peak Area) in the indicated *UV-treated* samples compared to the indicated (equivalent) *untreated* samples. Red/Orange boxes highlight the estimated 'Relative % Decrease' of Bpa-containing *mono*peptide for the Val252Bpa/Phe136Bpa-$TrwI_{B6}$ variants, respectively.

FIGURE EV3

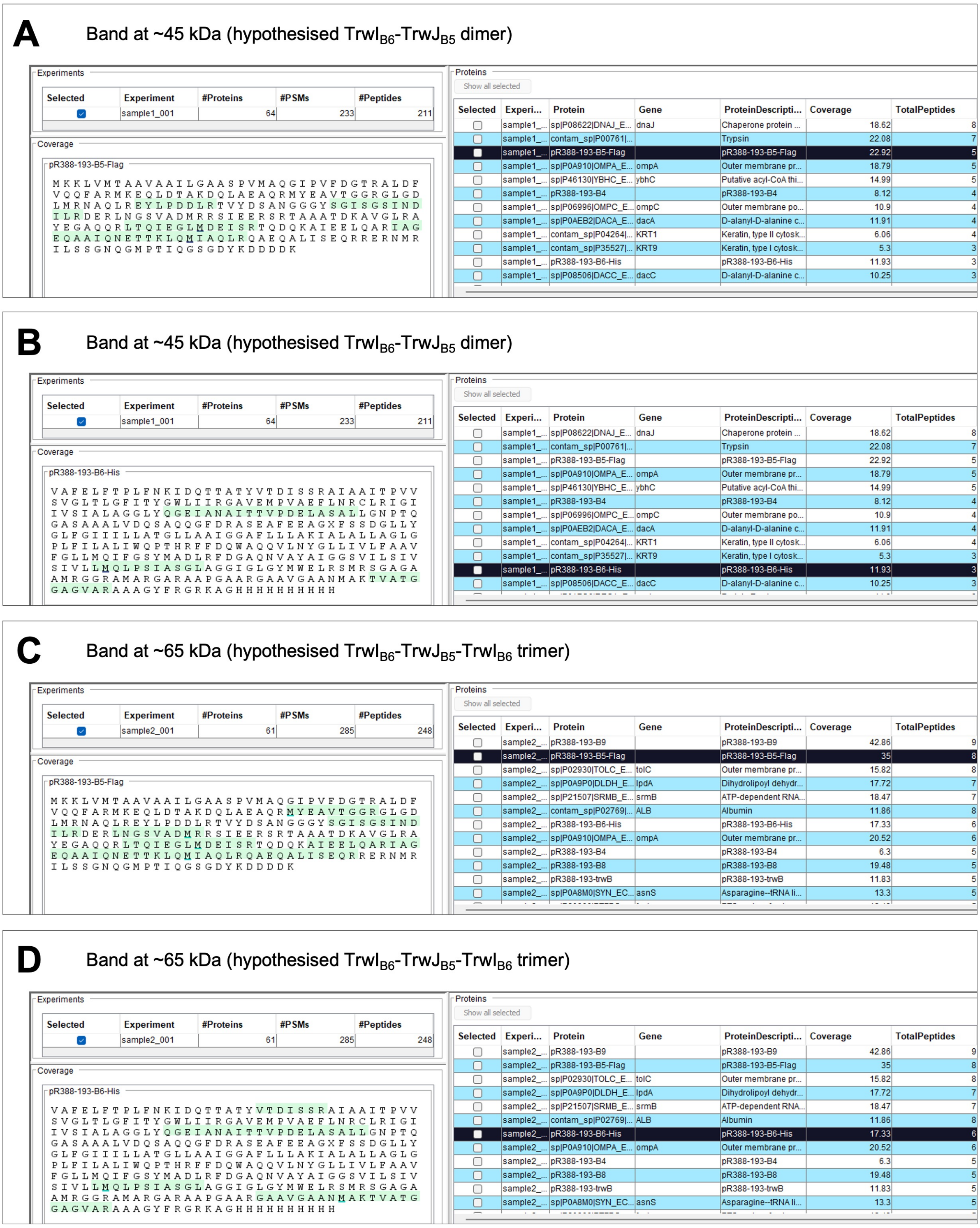
A
Band at ~45 kDa (hypothesised TrwI$_{B6}$-TrwJ$_{B5}$ dimer)
pR388-193-B5-Flag
B
Band at ~45 kDa (hypothesised TrwI$_{B6}$-TrwJ$_{B5}$ dimer)
pR388-193-B6-His
C
Band at ~65 kDa (hypothesised TrwI$_{B6}$-TrwJ$_{B5}$-TrwI$_{B6}$ trimer)
pR388-193-B5-Flag
D
Band at ~65 kDa (hypothesised TrwI$_{B6}$-TrwJ$_{B5}$-TrwI$_{B6}$ trimer)
pR388-193-B6-His

**Figure EV3**: *LC-MS/MS analyses confirming the presence of both TrwI$_{B6}$ and TrwJ$_{B5}$ in both gel bands*.
**A-D)** LC-MS/MS analysis of in-gel digestions of indicated gel bands (corresponding to either a hypothesised dimeric (~45 kDa) or trimeric (~65 kDa) species) from SDS-PAGE analysis of *in vivo* UV-treated, purified Phe136Bpa-TrwI$_{B6}$ variant (as shown in Fig 3F). Representation of relevant, theoretical, proteolytic peptides of TrwJ$_{B5}$ (A and C) and TrwI$_{B6}$ (B and D) in the datasets acquired from LC-MS/MS analyses of the trypsin/chymotrypsin in-gel digestions of the indicated samples were analysed and visualised using FragPipe (see Materials and Methods for details).

FIGURE EV4

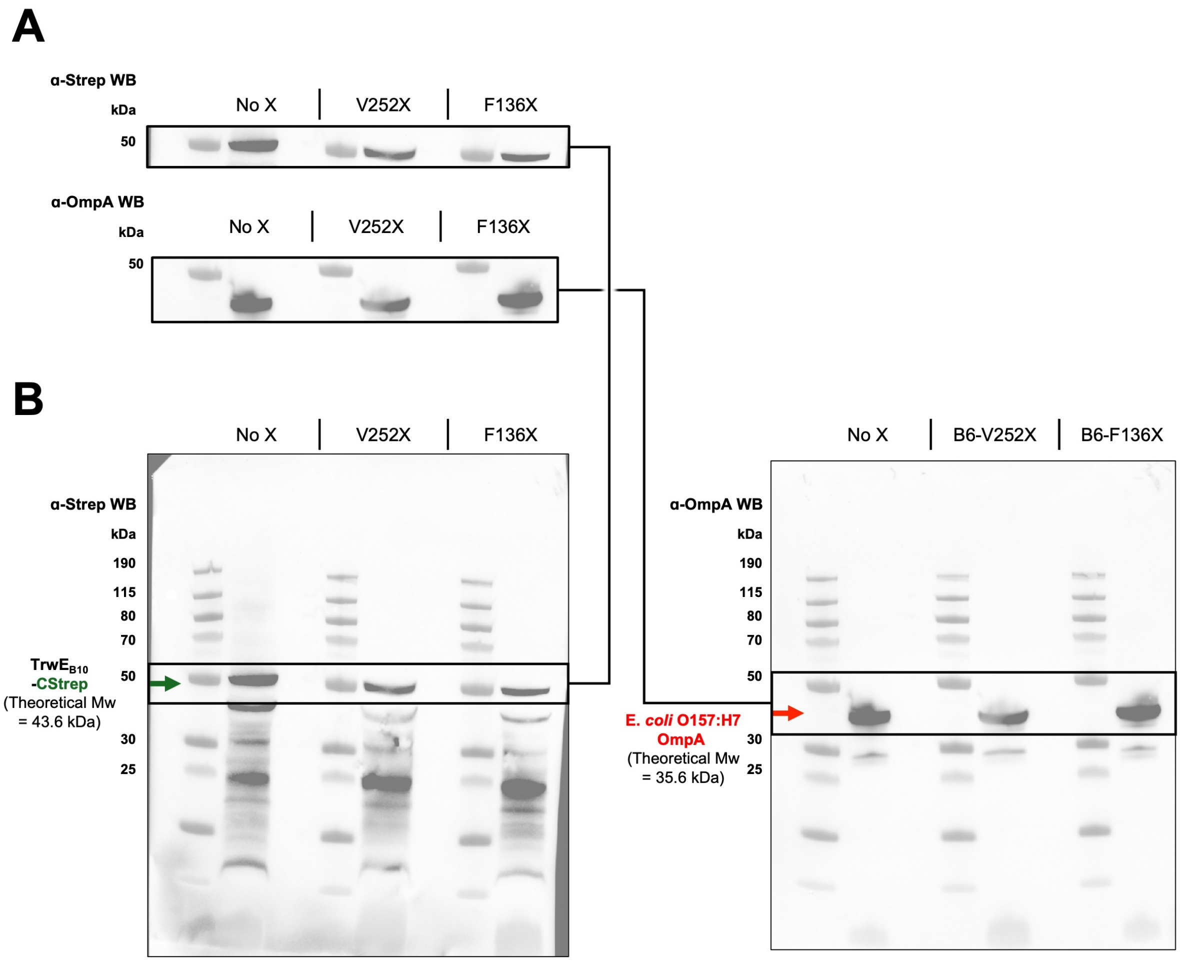
A
α-Strep WB
kDa
50
No X
V252X
F136X
α-OmpA WB
kDa
50
No X
V252X
F136X
B
No X
V252X
F136X
α-Strep WB
kDa
190
115
80
70
50
30
25
TrwEB10
-CStrep
(Theoretical Mw
= 43.6 kDa)
No X
B6-V252X
B6-F136X
α-OmpA WB
kDa
190
115
80
70
50
30
25
E. coli O157:H7
OmpA
(Theoretical Mw
= 35.6 kDa)

**Figure EV4**: *Western blot analysis confirming equal expression levels of complete R388 T4SS for each of the investigated donor cell variants*.
**A)** Selected sections of anti-Strep (top) and anti-OmpA (bottom) Western blots of whole cell extracts prepared from equal amounts (1 ODU, see Materials and Methods for details) of donor cell mass. The shown bands correspond to C-terminally Strep-tagged $TrwE_{B10}$ (used as proxy for estimation of relative quantity of complete T4SS present in the whole cell extract) and E. *coli* OmpA (loading control, used as proxy for estimation of total protein present in the analysed whole cell extract) (top and bottom panels, respectively).
**B)** Uncropped Western blots. The expected electrophoretic mobilities of C-terminally Strep-tagged $TrwE_{B10}$ and OmpA are indicated with coloured arrows (green and red, respectively).

FIGURE EV5

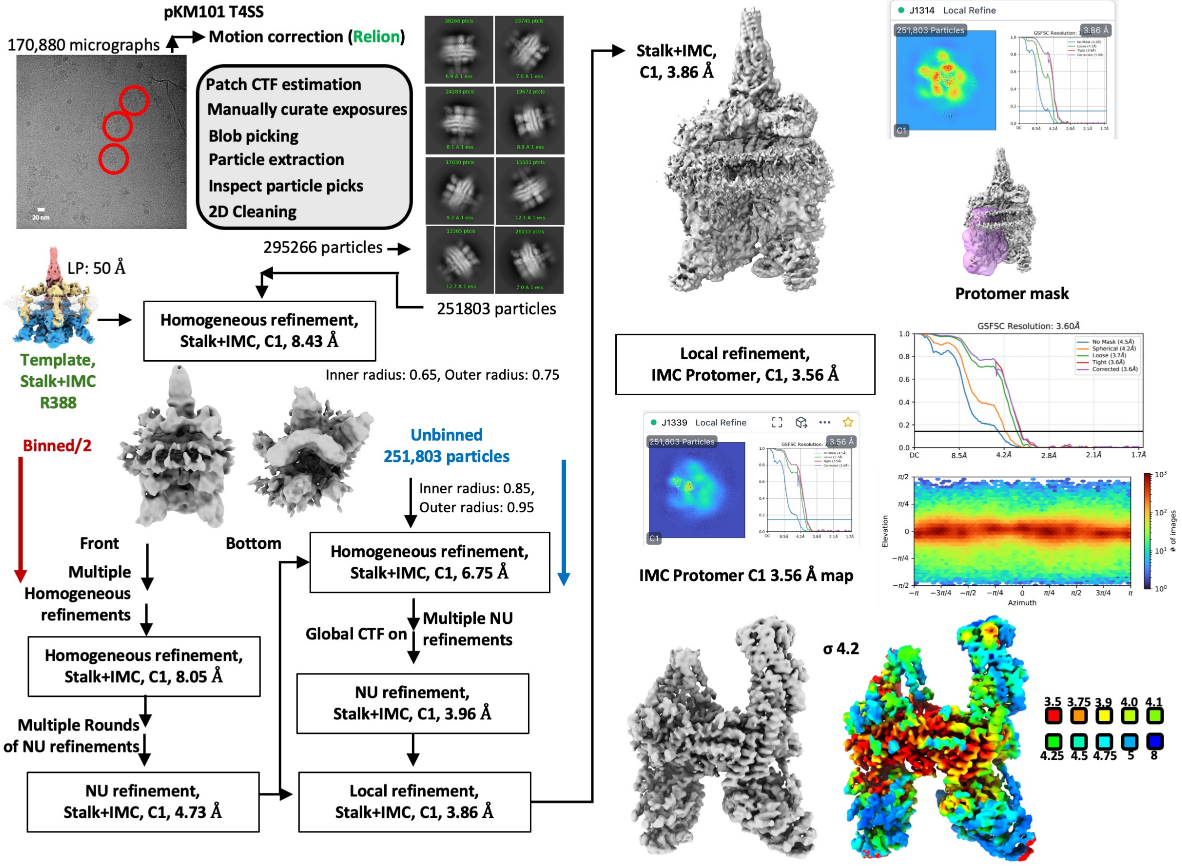
pKM101 T4SS
170,880 micrographs
Motion correction (Relion)
Patch CTF estimation
Manually curate exposures
Blob picking
Particle extraction
Inspect particle picks
2D Cleaning
295266 particles
251803 particles
LP: 50 Å
Template, Stalk+IMC R388
Homogeneous refinement, Stalk+IMC, C1, 8.43 Å
Inner radius: 0.65, Outer radius: 0.75
Binned/2
Front
Bottom
Unbinned 251,803 particles
Inner radius: 0.85, Outer radius: 0.95
Multiple Homogeneous refinements
Homogeneous refinement, Stalk+IMC, C1, 6.75 Å
Homogeneous refinement, Stalk+IMC, C1, 8.05 Å
Global CTF on
Multiple NU refinements
NU refinement, Stalk+IMC, C1, 3.96 Å
Multiple Rounds of NU refinements
NU refinement, Stalk+IMC, C1, 4.73 Å
Local refinement, Stalk+IMC, C1, 3.86 Å
Stalk+IMC, C1, 3.86 Å
J1314 Local Refine
251,803 Particles
Protomer mask
Local refinement, IMC Protomer, C1, 3.56 Å
GSFSC Resolution: 3.60Å
No Mask (4.5Å)
Spherical (4.2Å)
Loose (3.7Å)
Tight (3.6Å)
Corrected (3.6Å)
J1339 Local Refine
IMC Protomer C1 3.56 Å map
Elevation
Azimuth
### of images
σ 4.2
3.5
3.75
3.9
4.0
4.1
4.25
4.5
4.75
5
8

**Figure EV5**: *Overview of the cryo-EM data collection and processing that resulted in the final single particle reconstruction of the pKM101 IMC protomer.*
Representative micrograph (with scale bar) and representative 2D classes are shown, along with an overview of the data processing pipeline, featuring images of relevant maps, masks, FSC curves, and Euler angle distributions plots. The final Coulomb potential map of the IMC protomer is shown at a suitable contour level and coloured according to local resolution (in Å).

FIGURE EV6

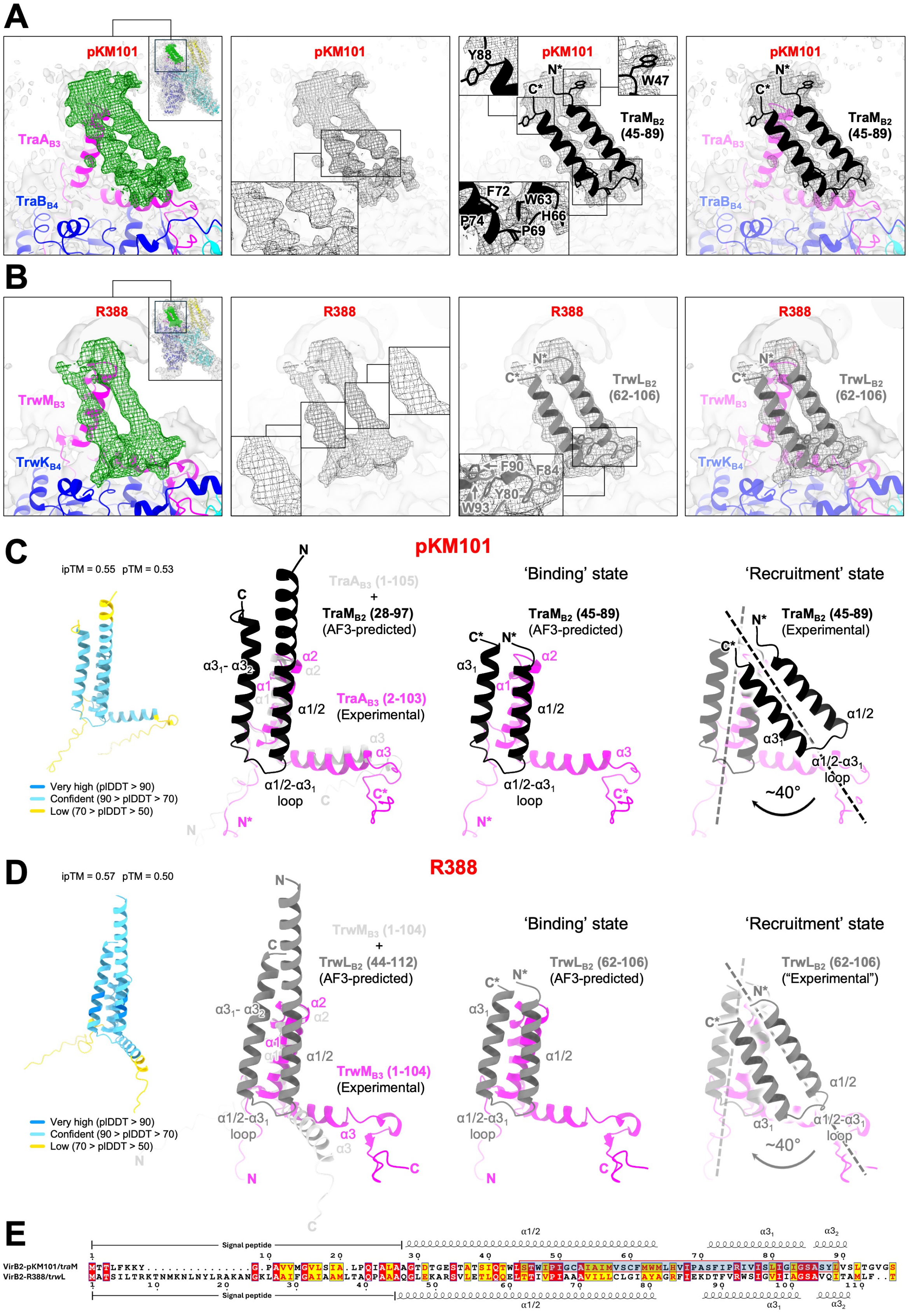

A
pKM101
TraA$_{B3}$
TraB$_{B4}$
Y88
N*
C*
W47
TraM$_{B2}$ (45-89)
F72
W63
H66
P74
P69
B
R388
TrwM$_{B3}$
TrwK$_{B4}$
TrwL$_{B2}$ (62-106)
F90
F84
Y80
W93
C
pKM101
ipTM = 0.55 pTM = 0.53
Very high (plDDT > 90)
Confident (90 > plDDT > 70)
Low (70 > plDDT > 50)
TraA$_{B3}$ (1-105)
+
TraM$_{B2}$ (28-97)
(AF3-predicted)
TraA$_{B3}$ (2-103)
(Experimental)
'Binding' state
TraM$_{B2}$ (45-89)
(AF3-predicted)
'Recruitment' state
TraM$_{B2}$ (45-89)
(Experimental)
~40°
D
R388
ipTM = 0.57 pTM = 0.50
TrwM$_{B3}$ (1-104)
+
TrwL$_{B2}$ (44-112)
(AF3-predicted)
TrwM$_{B3}$ (1-104)
(Experimental)
'Binding' state
TrwL$_{B2}$ (62-106)
(AF3-predicted)
'Recruitment' state
TrwL$_{B2}$ (62-106)
("Experimental")
~40°
E
Signal peptide
α1/2
VirB2-pKM101/traM
VirB2-R388/trwL

**Figure EV6**: *Cryo-EM and AF3-analysis of the pKM101 and R388 VirB2-VirB3 interaction*.
**A** and **B)** Visualisations of, respectively, the pKM101 and R388 IMC protomer cryo-EM maps (surface), featuring densities (mesh) into which models (cartoon and sticks) of, respectively, $TraM_{B2}$ and $TrwL_{B2}$ were fitted (see Materials and Methods for details). The C- and N-termini of the built residue stretches (45-89 for $TraM_{B2}$, 62-106 for $TrwL_{B2}$) are indicated, with the "*" indicating the termini of models in which not all residues of the mature protein are built. Selected density regions and model elements are highlighted (black boxes), and the complete IMC protomer maps (with fitted cartoon models) are shown as inserts.
**C** and **D)** Visualisations of AF3-predicted (AF3-predicted) and experimentally determined (Experimental) configurations of, respectively, the $TraM_{B2}$-$TraA_{B3}$ and $TrwL_{B2}$-$TrwM_{B3}$ interactions. The C- and N-termini of built residue stretches are indicated, with the "*" indicating the termini of models in which not all residues of the mature protein are built. The complete AF3-predictions of the binary VirB2-VirB3 complexes are shown, with individual residues coloured according to their pLDDT confidence score. The AF3-predicted VirB2-VirB3 structures are aligned to the VirB3 components (magenta) of the relevant IMC protomer models (see Materials and Methods for details), and the respective VirB2 moieties manually truncated to match the sequence stretches represented by distinct density in the relevant IMC protomer maps. Dashed lines (black) represent longitudinal axes of VirB2 in the (AF3-predicted) 'Binding state' and (experimentally determined = Experimental) 'Recruitment state', with curved arrows indicating the pivotal motion of VirB2 apparently relating the states.
**E)** Amino acid sequence alignment of pKM101 $TraM_{B2}$ and R388 $TrwL_{B2}$ with select sequence elements indicated above and below, respectively. Residues are coloured according to sequence homology (red = identity, yellow = conservation), and the sequence stretch of $TraM_{B2}$ successfully built into the cryo-EM map of the pKM101 IMC protomer (see Materials and Methods for details) is highlighted in blue.

#### FIGURE EV7

### A

**Relative TrwL/VirB2 positioning** when respective pairs of incorporated TrwL/VirB2 + TrwM/VirB3 are aligned along TrwM/VirB3
Cα RMSD (TrwL/VirB2 : TrwL/VirB2)

| Cα RMSD (Å) | Pos 1 / Cycle X+1 | Pos 2 / Cycle X+2 | Pos 3 / Cycle X+3 | Pos 4 / Cycle X+4 | Pos 5 / Cycle X+5 | Pos 6 / Cycle X+6 |
|---|---|---|---|---|---|---|
| Pos 1 / Cycle X+1 | 0.000 | 1.454 | 2.265 | 2.366 | 1.799 | 0.568 |

### B

Pos 1 / Cycle X+1

Pos 2 / Cycle X+2

Pos 2 / Cycle X+2
rigid body aligned to
Pos 1

Cα-RMSD
TrwL/VirB2 Cycle X+2
vs.
TrwL/VirB2 Cycle X+1
= 1.454 Å

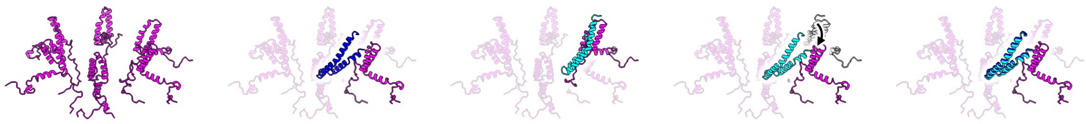

### C

**Relative TrwL/VirB2 positioning** (at incorporation stage)
Cα RMSD (TrwL/VirB2 : TrwL/VirB2)

| Cα RMSD (Å) | Pos 1 / Cycle X+1 | Pos 1 / Cycle X+7 | Pos 1 / Cycle X+13 | Pos 1 / Cycle X+19 | Pos 1 / Cycle X+25 | Pos 1 / Cycle X+31 | ... | Pos 1 / Cycle X+55 |
|---|---|---|---|---|---|---|---|---|
| Pos 1 / Cycle X+1 | 0.000 | 0.789 | 1.176 | 1.296 | 1.266 | 1.264 | ... | 1.652 |

### D

Pos 1 / Cycle X+1

Pos 2 / Cycle X+55

Pos 2 / Cycle X+1
+
Pos 2 / Cycle X+55

Cα-RMSD
TrwL/VirB2 Cycle X+1
vs.
TrwL/VirB2 Cycle X+55
= 1.652 Å

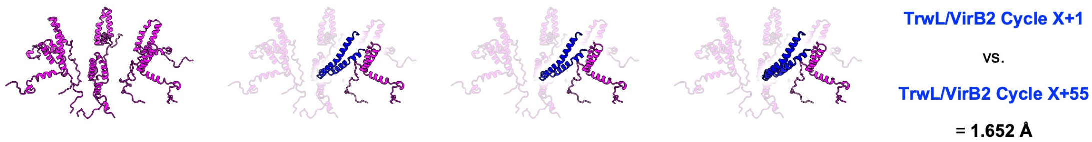

**Figure EV7**: *Demonstration of geometric consistency of the proposed model.*
**A)** Table listing Cα RMSD values (in Å) between the $TrwL_{B2}$ moieties of the indicated pairs of $TrwL_{B2}$ and $TrwM_{B3}$ (the pairs are treated as independent models) at the end of the indicated incorporation cycles. The individual pairs have been rigid body aligned along their respective $TrwM_{B3}$ moieties. The slight deviations reflect the fact that every $TrwL_{B2}$ monomer within the pilus structure was considered as an independent entity during model building (Vadakkepat *et al*, 2024) with no helical symmetry or symmetric packing arrangements being imposed. As is evident, these slight deviations systematically fall within expected levels of modelling uncertainty.
**B)** Visual representation of how the values listed in panel A) were obtained. All modelling and calculations were done using ChimeraX (see Materials and Methods for details).
**C)** Same as in panel A), except no rigid body alignment (only superimposition) was performed. These calculations thus reflect that consistent (i.e. within expected levels of modelling uncertainty) accommodation of the sixfold/fivefold IMC/pilus symmetry mismatch is achieved by letting each incorporation cycle culminate in a ~6°/~2.6Å rotation/rise of the entire nascent pilus.
**D)** Visual representation of how the values listed in panel A) were obtained. The modelling and calculations for the state of pilus biogenesis shown in Fig 7E (end of incorporation cycle X + 55) are shown.
